# Cultivation of *Ulva* sp. offshore the Eastern Mediterranean Sea in experimental bioreactors: seasonal growth dynamics and environmental effects

**DOI:** 10.1101/2023.01.15.524102

**Authors:** Meiron Zollmann, Alexander Liberzon, Ruslana R. Palatnik, David Zilberman, Alexander Golberg

## Abstract

Offshore macroalgae production could provide an alternative source of biomass for food, materials and energy. However, the offshore environment in general, and specifically the Eastern Mediterranean Sea (EMS) offshore, is a high energy and low nutrients environment and thus is challenging for macroalgae farming. This study aims to understand the effects of season, depth, and fertilization duration on growth rates and chemical composition in offshore *Ulva* biomass production and develop a predictive model suitable to offshore conditions. We hypothesize that offshore *Ulva* growth rates and chemical composition will follow a seasonal trend and that applying rapid onshore fertilization could refill nutrient storages and enable continuous offshore cultivation. We test this hypothesis by measuring *Ulva* biomass and internal nitrogen in offshore experiments in the nitrogen-poor EMS a few kilometers offshore the Israeli coast. We construct a predictive cultivation model to estimate N concentrations in the sea during experiments. This study demonstrates the feasibility of growing *Ulva* sp. offshore the EMS with an onshore nutrient supply and develops a better understanding of seasonal growth dynamics and environmental effects (nitrogen, waves, depth, etc.). Furthermore, the study showcases the applicability of the macroalgae cultivation model in the offshore environment and its potential contribution throughout the whole lifecycle of seaweed cultivation.

## 1. Introduction

Increasing global demand for food, chemicals, materials and fuel increases the demand for crops and biomass, thus, increasing pressure on the agricultural systems. Offshore-grown macroalgae could provide an alternative source of biomass without any need for arable land or fresh water. However, macroalgae still present only a tiny percent of the global biomass supply of ∼17·10^6^ ton Fresh Weight (F.W.) of macroalgae compared to 16·10^11^ tons of terrestrial crops, grasses, and forests ^1–3^. An expanding body of evidence suggests that offshore macroalgae biomass cultivation, rather than harvesting wildstock, can provide biomass for the sustainable production of food, chemicals, and fuels ^1,4,5^.

The global potential of offshore biorefineries to provide for biomass for proteins, platform chemicals, and energy was analyzed by ^6^, using a metabolism model of the green marine macroalga *Ulva*, coupled with essential inputs from climatological oceanographic data. *Ulva* is an interesting biomass for biorefinery as it grows globally and it was already converted to protein, cellulose, starch, ulvan (bioactive polisacharide), fatty acids, minerals, biocrude, biochar, ethanol, biogas^7^, and polyhydroalkanolyes (PHA)^8^, all of which could serve as building blocks for a sustainable bioeconomy^9^.

However, production and processing of the biomass, in the offshore high energy and low nutrients environments is challenging. The current offshore marine biomass cultivation concepts include Near Farm Concepts for kelp growth^10^, Tidal Flat Farms, Floating Cultivation^10^, Ring Cultivation^11^, wind-farm integrated systems^12^, and air mixed cages^13^. Artificial fertilizing is generally not recommended but if needed, can be performed via integration to fish farms in an Integrated Multitrophic Aquaculture (IMTA)^14^ or artificial upwelling of nutrient-rich deep water^15^. Another potential fertilizing solution, although logistically challenging, to utilize Ulva’s high uptake rates (up to 470 μmol N g^-1^ D.W. h^-1^) ^16–20^ and recharge critical nutrients by rapid fertilizing in a designated fertilizing tank, which can be performed on or offshore. Due to the high complexity and sophisticated work logistics in the offshore environment, establishing large offshore farms for biomass production requires rigorous planning and attention to various design and operation parameters. Parameters such as cultivation depth, cultivation cycle, and pre-cultivation fertilization should be determined based on understanding seasonal growth dynamics and environmental effects and are critical for economically viable production.

Experience from terrestrial agriculture shows that dynamic models that aim to predict the field yield significantly improve the economics and long-term sustainability of food supply systems^21^. Such detailed models, which combine biomass productivity, crop yields, sustainability, and economics, are not yet available for macroalgae offshore farms. Yet, initial steps in this direction have been made by developing growth function models to predict seasonal blooms ^22–33^ and biomass productivity in the controlled photobioreactors ^34–38^. The existing models for offshore biomass productivity do not provide information on the effects of dynamic environmental factors such as light, temperature and nutrients on the biomass productivity and chemical composition. Thus, their applicability for the design of the real-time seaweed farms is low ^39,40^.

This study aims to understand the effects of seasonality, depth, and fertilization duration on growth rates and chemical composition in offshore *Ulva* biomass production and develop a predictive model suitable to offshore conditions. We investigated the variation of *Ulva* biomass productivity and internal nitrogen in the offshore experiments in nitrogen poor (oligotrophic) Eastern Mediterranean Sea (EMS) a few kilometers offshore the Israeli coast. We constructed a predictive cultivation model based on previous models, which were adjusted to the offshore conditions. We tested its performance using experimental data from an offshore experimental station tailor designed for this study.

## 2. Results

The results comprise of a) experimental results of growth rates and chemical compositions of *Ulva* sp. macroalgae cultivated in cages offshore the EMS under different conditions, and b) model simulations for *Ulva* sp. growth rates and chemical compositions. *Ulva* sp. cultivation experiments in offshore cages provide year-round growth rates and chemical compositions in the Israeli EMS in two depths (1 and 5 m). In addition, it examines the effectiveness of short-term fertilizing treatments between two low-nutrient cultivation periods. All experiments followed a similar design, validating the cultivation model under naturally varying environmental conditions offshore (Section 2.4). DGR and internal N data are summarized in Table S7 and Table S8 (Appendix F). Data analysis was performed in a step-by-step manner, starting with the effect of pre-cultivation fertilization regime (continuous vs. rapid, Section 2.1), continuing to the effect of cultivation depth (Section 2.2), and finishing with the effect of cultivation date (Section 2.3).

### 2.1 Fertilizing regime

One day (rapid) onshore fertilizing method effectivity is compared to that of continuous nutrient enrichment method using the DGR and internal N. Results from experiment #1 show higher DGR (p-value < 0.05) and slightly higher internal N for the continuous fertilization (Figure 1). Figure S4 signifies the result comparing the results of each depth (1 and 5 meters cages) separately (p-value < 0.001), whereas there is a consistent but relatively mild effect on internal N in both depths. In the rapid fertilizing experiments, no difference can be found between DGR of samples fertilized in regular seawater and samples fertilized in enriched seawater. Based on these results, we decided to focus on the experiments performed after continuous nutrient enrichment in the MPBR, as opposed to the rapid onshore fertilizing that was proven ineffective in these conditions.

**Figure 1.**
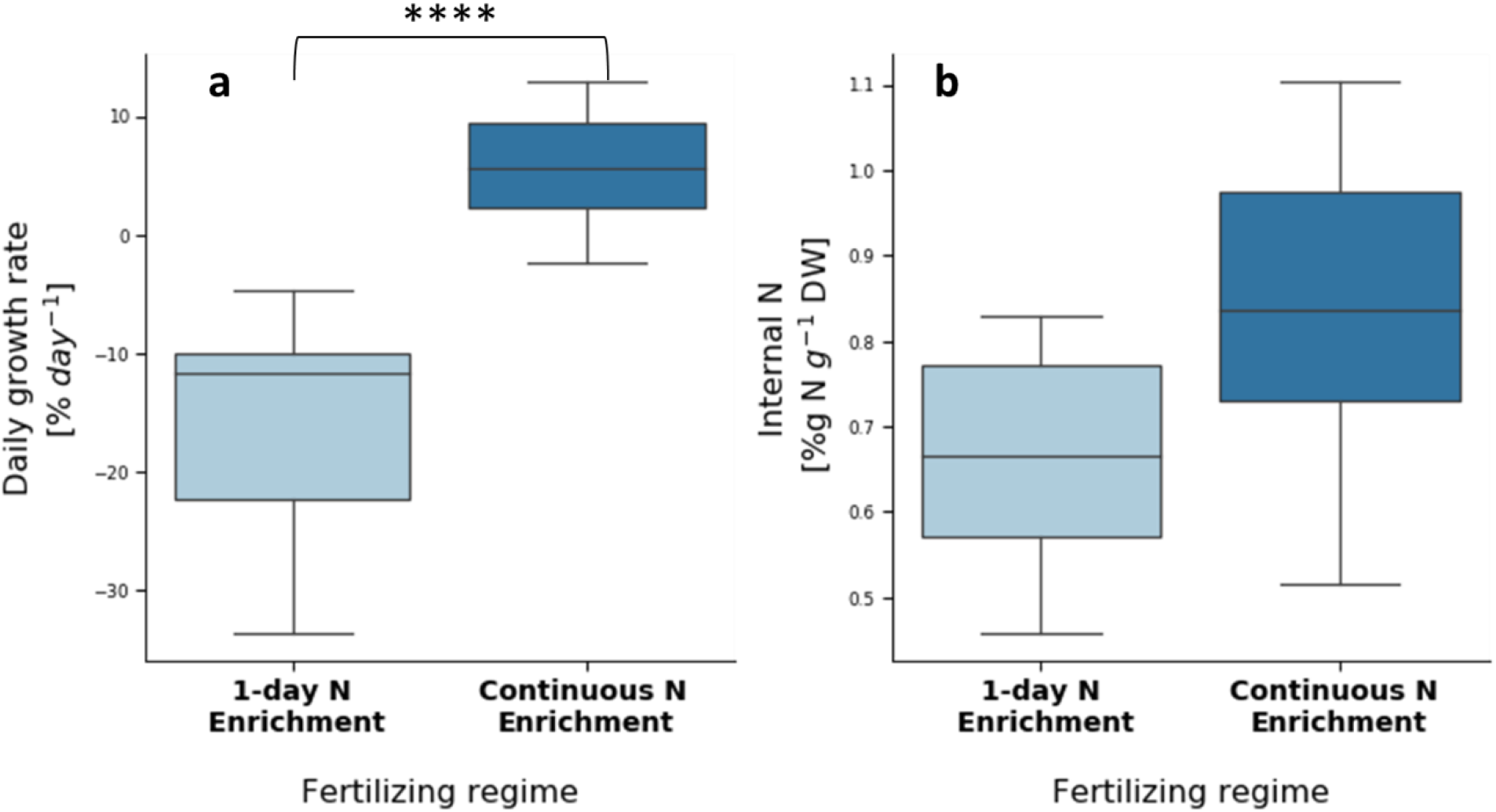
Daily growth rate **(a)** and internal N **(b)** of *Ulva* sp. cultivated offshore the EMS after rapid nutrient enrichment **(light blue)** and after continuous nutrient enrichment **(dark blue)**. Asterisks indicate statistical significance of difference with **** p < 0.0001, calculated by the two-tailed Mann-Whitney U test. # samples: rapid nutrient enrichment: 22 for DGR and 7 for internal N. continuous nutrient enrichment: 54 for DGR and 21 for internal N.

### 2.2 Cultivation depth

Cultivation of *Ulva* sp. offshore the EMS in a depth of 5 m has shown higher growth rates (p-value < 0.0001, N = 30) than cultivation in a depth of 1 m, but similar internal N values (N = 8). We compare (Figure 2) the results of experiment #1 and experiment #2, with cultivations at both depths. We attribute the results to different environmental conditions, specifically waves and currents. Visual observations (Figure S5 and Figure S6) point out an important difference between depths. Whereas in the 5m cages the biomass is distributed throughout the whole surface of the cages, in the 1m cages the biomass clumped on the cage frame. This distribution difference indicates that the water motion, produced by waves and currents, is more prominent in the 1 m depth. Based on this observation, we suggest two potential mechanisms causing lower growth rates in the 1m depth: 1. Faster mechanical wearing and biomass losses in the shallower depth, and/or 2. biomass clumping inhibited growth due to a smaller surface area and a lower exposure to light.

**Figure 2.**
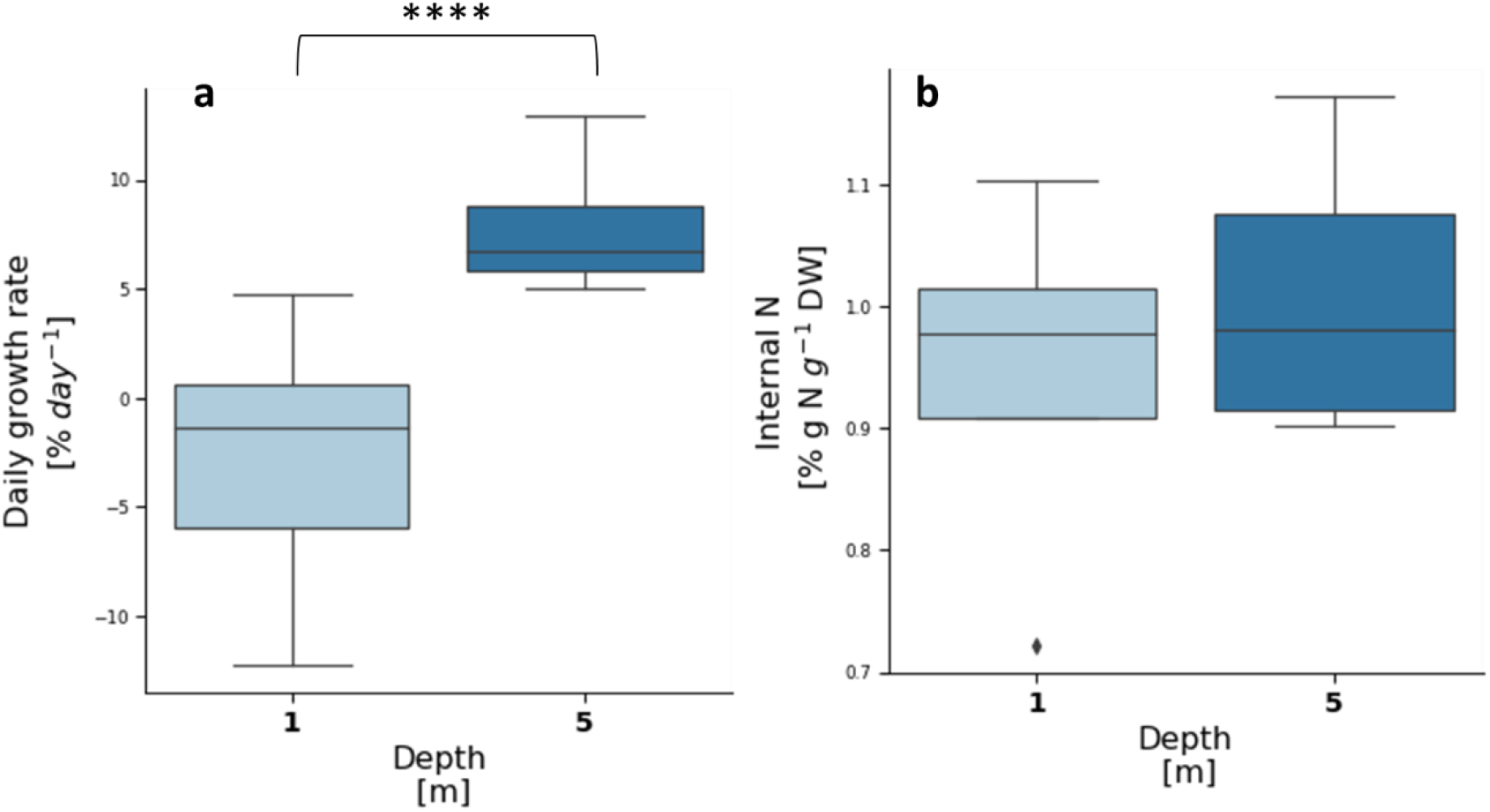
Daily growth rates **(a)** and internal N **(b)** of *Ulva* sp. cultivated offshore the EMS after continuous nutrient enrichment in depths of 1 m **(light blue)** and 5 m **(dark blue)** in experiments 1 and 2. Asterisks indicate statistical significance of difference with **** p < 0.0001, calculated by the two-tailed Mann-Whitney U test. # samples: 15 for DGR and 4 for internal N for each depth.

### 2.3 Timeline of offshore experiments and seasonal effects

We compare here results from four experiments from only 5 m depth experiments with continuous fertilization method. Visual observation (see harvested cages in Figure S7) supports the results that growth rates vary seasonally (Figure 3, left) (p-value < 0.0001). Multiple comparison tests found that growth rates in December 2019 were significantly lower (and in some cases negative) than all other experiments that are comparable in DGR (p-value < 0.05).

**Figure 3.**
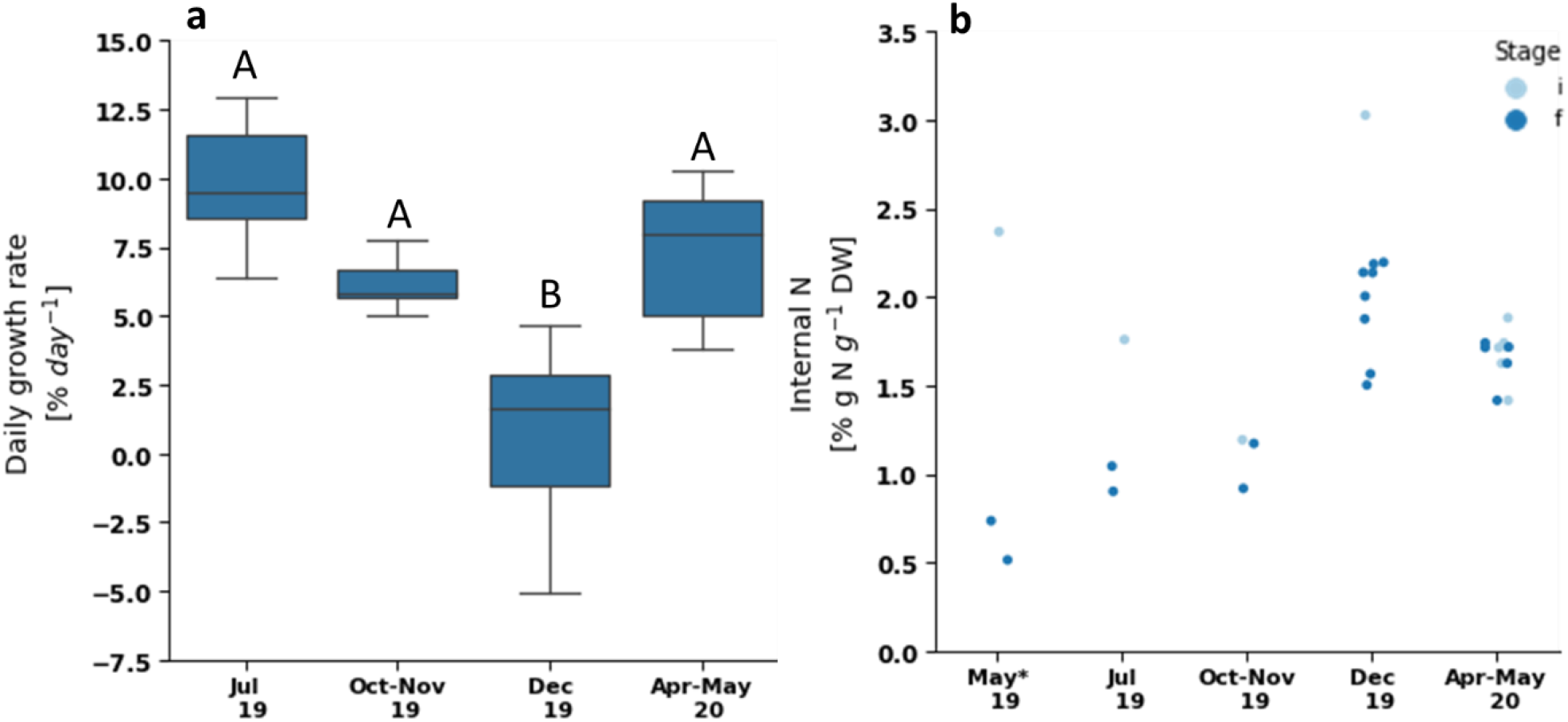
Daily growth rate **(a)** and internal N **(b)** of *Ulva* sp. cultivated offshore the EMS after continuous nutrient enrichment in a depth of 5 meters, grouped by experiment date. Group A is statistically different from group B, in a significance of p < 0.05, calculated by the post-hoc Dunn’s test with the Bonferroni adjustment method for pairwise comparison. On the right, red dots present internal N at the beginning of each experiment and green dots represent internal N at the end of each experiment. # samples: 6-12 for DGR and 2-8 for internal N for the different experiments. *From the preliminary run (May 19) we show only internal N results, as the DGR measurements are meaningless due to biomass losses.

We study weather conditions that led the DGR and internal N at different experiments, specifically irradiance, rain, waves, and currents (Appendix I). The low growth in the December experiments due to some combination of stormy weather: rough sea conditions and light limitation. Figure S9 and a report of the staff of the adjacent Lev-Yam fish cages facility (personal communication) present that the first few days of the December cultivation period were stormy, with a significant wave height (highest third of the waves) higher than 2.5 m. Four cages show biomass losses, probably a result of the waves. The biomass at 5 m depth is also clumped on the cage frame in these conditions (Figure S7, second row), similar to the 1 m cages presented above (Figure S5 and Figure S6). The stormy days induce also light limitation through two different mechanisms: clouds reduce global irradiance at the sea surface (see Figure S8) and significant rain event on December 12 (28 mm per day, based on IMS data), that typically causes flash floods in the Alexander estuary, resulting in coastal enrichment of particulate matter and an increased turbidity. It is noteworthy that during the April-May cultivation period, the significant wave height was relatively high (around 2 m) and some rainfall was measured (5 mm during 4-5 May), but growth rate was high. A prominent difference between different high-wave periods is in the wind direction: western winds during December versus Eastern winds during May. Although it is impossible to pinpoint the dominant mechanism, we believe that the high turbidity and strong waves are the equally probable. In the winter experiment (December 19), half of the cages were harvested right after the storm and half remained offshore for a longer period. The cages harvested right after the storm had larger portion of biomass loss (three cages versus 1) compared to the cages that remained for another week, pointing to the biomass recovery during the additional five days. Higher irradiance (i.e., less clouds and lower turbidity) and lower waves enabled the biomass to grow and accumulate despite initial losses (see cages in Figure S7, third row from the top). Internal N in the longer cultivation duration slightly decreased, consistent with the growth rates in a nutrient poor environment. We conclude that winter growth is possible with longer cultivation periods after storms and rainfall, as long as the nutrients are provided to keep the internal N around 1 % g N g^-1^ D.W. (Figure 3, right).

Internal N decreases during the cultivation experiments in respect to the initial internal N (taking into account also the results of the preliminary experiment) (Figure 3, right, p-value < 0.01). Higher DGR leads to lower internal N in experiments #1-4 (Pearson’s r = -0.462) supporting the growth model hypothesis that in an oligotrophic environment growth is based mostly on initial internal N. However, there are unusually high internal N results in experiments #2 and #4 in respect to the high DGR, suggesting that there was an external additional nutrient supply to the cultivation site during these periods. This suggestion is supported also by the mass balance presented in Table 1, confirming that N uptake was substantial in those two periods, especially during April-May 2020. We attribute this supply to the sea currents from the rich nutrient site of Alexander estuary (see Section 3 for further discussion).

**Table 1.**
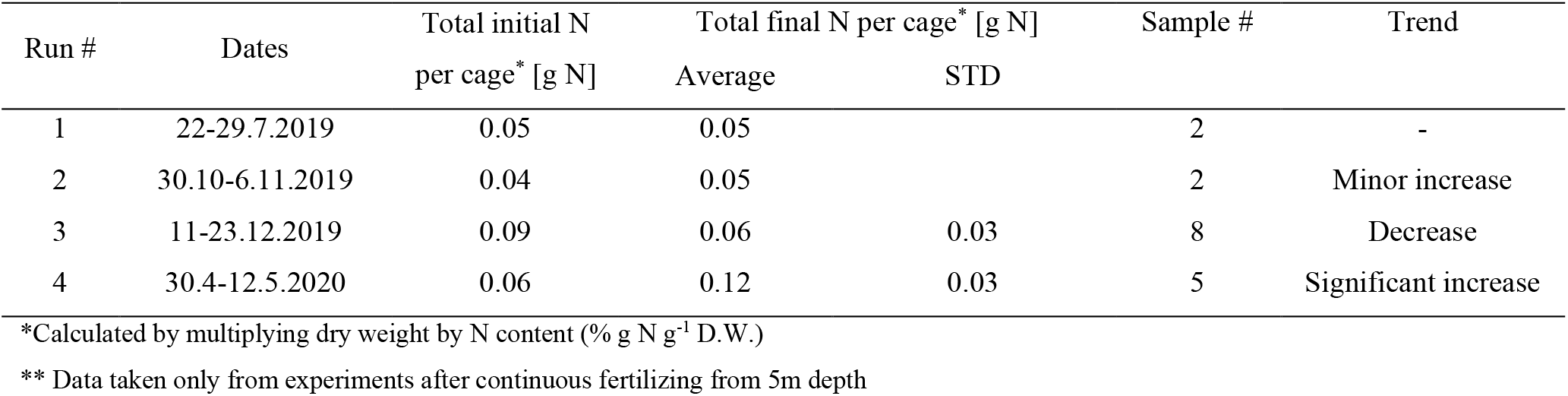
Nitrogen mass balance in offshore experiments.

| Run # | Dates | Total initial N | Total final N per cage* [g N] |  | Sample # | Trend |
| --- | --- | --- | --- | --- | --- | --- |
|  |  | per cage* [g N] | Average | STD |  |  |
| 1 | 22-29.7.2019 | 0.05 | 0.05 |  | 2 | - |
| 2 | 30.10-6.11.2019 | 0.04 | 0.05 |  | 2 | Minor increase |
| 3 | 11-23.12.2019 | 0.09 | 0.06 | 0.03 | 8 | Decrease |
| 4 | 30.4-12.5.2020 | 0.06 | 0.12 | 0.03 | 5 | Significant increase |

\*Calculated by multiplying dry weight by N content (% g N g<sup>-1</sup> D.W.)
\*\* Data taken only from experiments after continuous fertilizing from 5m depth

### 2.4 Offshore cultivation simulations

The main purpose of the mathematical model and the associated software (see Supplementary Materials) is to estimate the external N supply to the offshore cultivation sites and predict growth rates and internal N conditions for optimal cultivation cycles.

The analysis is of the results of cultivation in a depth of 5 m after continuous fertilization in experiments #1, 2, and 4. Experiment #3 (December 2019) is excluded from the model due to a storm related biomass loss.

Additional factors that we considered are the current regime, potentially changing external N concentration, and the wave height, affecting biomass distribution inside the cages. To alleviate the effect of unequal biomass distribution (Figure S7, bottom row) after the high wave event experienced on May 6 (Table 2 and Figure S9), we assumed that the surface area covered by *Ulva* decreased by a factor of 2 after the high waves. Considering both factors, we assumed that the initial N concentration in the sea was similar to the background concentration when the dominant current direction was North-Easterly (i.e. 0.75 μM N, as was estimated for experiment #1), and that it increased following the change of current direction from Northly to Westerly.

**Table 2.** Sea conditions, model errors and estimated offshore N concentrations during the offshore experiments.

| Experiment # | Dates | Dominant Currents | Waves [m] | Estimated $N_{ext}$ [$\mu\text{M N}$] | $m$ # samples | $N_{int}$ # samples | Mean RMSRE [%] |
| --- | --- | --- | --- | --- | --- | --- | --- |
| 1 | 22-29.7.2019 | North-Easterly | < 1 | 0.75 | 5 | 2 | 27.1 |
| 2 | 30.10-6.11.2019 | Westerly | < 1 | 1.25 | 9 | 2 | 18.1 |
| 4 | 30.4-12.5.2020 | 30.4-7.5: North-Easterly<br>8-12.5: Westerly | 30.4-5.5: < 1<br>6.5: ~2.5<br>7-12.5: < 1.5 | 30.4-7.5: 0.75<br>8-12.5: 3.6 | 12 | 5 | 42.8 |

Simulation results leading to the estimate of *N*_*ext*_, are presented below in Figure 4: experiment #1 in the left column, experiment #2 in the middle column and experiment #4 in the right column. In experiments #1 and #2 the simulation predicts the outcome of the DGR and internal N with satisfactory accuracy (RMSRE = 27.1 or 18.1 %). In experiment #4 we observe a larger error (RMSRE = 45.1 %) that we attribute to the incorrect settings of N_ext_. Artificially adding external N at the time instant after the storm, when the current direction changed, is helpful from the simulation point of view and improves the obtained result (RMSRE = 42.8 %). Other modifications, e.g. irradiance or temperature cannot explain the results at the same efficiency.

**Figure 4.**
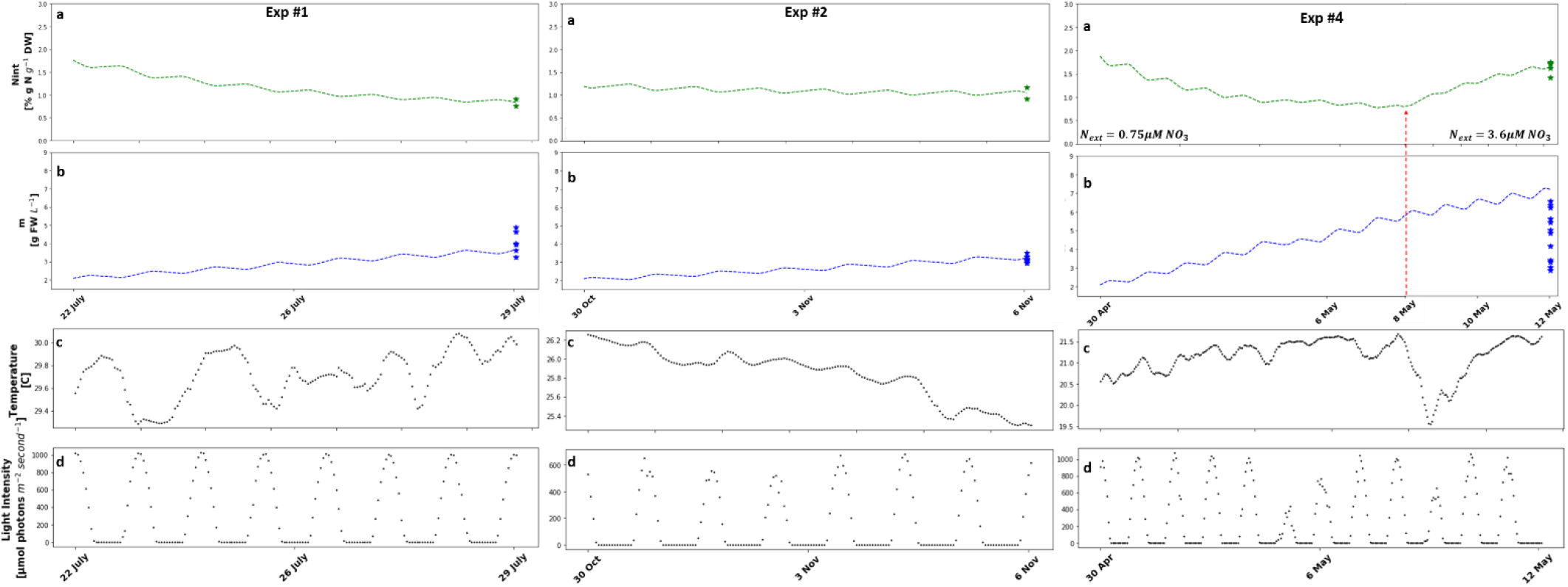
Model timewise simulation of *Ulva* sp. cultivation in offshore cages for a period of 7-10 days in experiments #1 (**left**), #2 (**center**) and #4 (**right**). Two variables are followed: *N*_*int*_ (% g N g^-1^ DW, **a**) and *m* (g DW L^-1^, **b**). Biomass initial conditions: 2.08 g D.W. per L^-1^ (20 g F.W. per cage). *N*_*int*_ initial conditions: 1.76, 1.19 and 1.88 % g N g^-1^ D.W. for experiments #1, #2 and #4, respectively. *N*_*ext*_ initial conditions: 0.75 and 1.25 μM N for experiments #1 and #2, respectively. In experiment #4 N concentration in seawater changes from 0.75 μM N initially to ∼3.6 μM N after the May 8 (marked by red arrow). Empiric data points are presented in stars. (**c**) and (**d**) Temperature and light intensity profiles.

We take the simulation results with caution. The sensitivity analysis of the model shows that it is sensitive to *K*_*a*_ and *K*_*I*_ (0.3-0.5, Figure S13). We could suggest that large biomass density in these unmixed cages leads to smaller light penetration depth and as a result, increased internal N storage. This does not diminish the dependence of internal N model on *K*_*s*_ and *V*_*max*_ (0.2-0.55), both pointing to dominant effect of the external N concentration in the environment. Biomass production in DGR is more sensitive to temperature related parameters, *T*_*opt*_, *T*_*max*_ and *n* (0.1-0.2), such that temperature rise to around 30°C during the summer can be set as a limiting, or maximum possible growth temperature for *Ulva* sp. ^47^. Future studies could focus on additional effects of light absorption and that of active mixing of *Ulva* sp. biomass in offshore cages ^13^.

In conclusion, we presented how with some minor adjustments, the model that was developed for a controlled mixed reactor can be used to simulate the dynamics of biomass production and internal N also in an offshore environment and a flat cage cultivation system.

Furthermore, we developed a method to estimate N level in the sea during the cultivation period. This method should be validated by actual measurements of N concentrations (nitrate and ammonium) in the sea during an offshore experiment.

## 3. Discussion

This study demonstrates the feasibility of offshore cultivation of *Ulva* sp. in the Eastern Mediterranean Sea and emphasizes the dominant effects: seasonal and environmental conditions, a necessary depth, and onshore fertilization in the absence of a sustainable offshore external N concentration. Furthermore, a supplementary model adds predictability to offshore cultivation, allowing for the optimization of cultivation cycles and fertilization requirements. The lowest growth rates offshore *Ulva* sp. were measured in the winter (December) and maximal growth rates in spring (May) and summer (July) (Figure 3). Although growth is possible during winter, the accumulated growth rates are low due to biomass losses during winter storms, turbidity, and strong currents. The 5 m depth is more favorable for *Ulva* sp. cultivation than the 1m depth. The inferiority of the 1m depth in terms of lower growth rates (Figure 2) is strongly associated with the mechanical stress caused by the surface waves (see cages in Figure S5 and Figure S6). Fertilization significantly affects growth rates, mainly because the existing external N concentration in the EMS is insufficient for sustained growth. The high peak, short-term (20-24 hours) fertilization treatment was ineffective in the examined conditions. This is in contrast to the model prediction (Figure S14) that 20 hours of fertilization can lead to the recovery of more than 1 % g N g^-1^ D.W. and support significant growth in the following week.

The idea of onshore preparation of the seaweed before offshore cultivation is not new^48^. Traditionally, seaweed spores and seedlings are maintained, propagated, and seeded on the cultivation rope/net and sometimes even fertilized^48^ in an onshore nursery, but a major part of the growth occurs in the sea, based on near/offshore nutrient concentrations. In an oligotrophic environment such as the EMS, however, growth will usually be limited by nutrient supply. Thus, onshore fertilizing and the initial N level at the beginning of the offshore cultivation period is essential. Here, we develop a framework for using cultivation models to optimize the operation of this two-stage cultivation method. An offshore cultivation model can predict growth and decide upon harvest timing. An onshore pond cultivation model can be used to design the onshore fertilizing stage according to the required biomass and level of internal N for the offshore stage. In addition, the onshore stage can be used to fine-tune model parameters in different seasons, locations, or seaweed species for improved on and offshore model performance. Although harder to calibrate, the offshore model can develop into an important tool for governments and seaweed farmers, providing seasonal predictions that will enable better preparation and operation in an era of increasing climatic uncertainty (i.e., predict how sea water temperature rise will affect the yields).

In experiments #2 and #4, a net N uptake was measured, implying that environmental N levels were not as low as previously reported^49^. Unfortunately, due to technical limitations, ammonia quantification in water samples was performed only during the preliminary experiments and experiments #1-#2, constraining our ability to support this also with water analysis data. Nevertheless, ammonia measurements demonstrated values in the range of 0-1.5 μM NH_4_ and did not show a consistent seasonal trend. These values fit the models *N*_*ext*_ estimations of 0.75-1.25 μM but are a bit lower than the 2 μM NH_4_ background values measured at a nearby site during the summer and autumn of 2012 in a study by Korzen et. al^49^.

The cultivation model explains the results, demonstrating that offshore N levels changed between experiments. The influx with the highest concentration was estimated in May 2020. In two offshore cultivation experiments, from November 2019 and May 2020, extra N uptake was recognized compared to the uptake expected according to background N_ext_ concentrations and model predictions. This extra N uptake could be associated with nitrogen effluents from the nearby Lev-Yam fish cages or with anthropogenic nutrients discharged from the Alexander estuary. Based on an assessment of dynamic current regimes (Appendix I) and constant fish loads (personal communication) in the fish cages, we estimate that the source of these elevated N levels is coastal, namely from the Alexander estuary.

Finally, we believe that this work sheds light on the seasonal growth dynamics and major environmental effects on *Ulva* sp. cultivation offshore the EMS. Our work emphasizes the importance of sufficient nutrient supply and the significant effect of surface water motion on offshore biomass yields. Notwithstanding, the presented analysis is limited in the scale, duration of the offshore experiments and the resolution of biomass and water sampling. Furthermore, the proposed model, despite its potential contribution to the progress of the field, will require additional data to improve its predictive abilities. In addition, the usability of the model for farm management and farm scale prediction may be elevated by incorporating it into a multi-scale model^39^.

## 4. Conclusions

We demonstrated the feasibility of growing *Ulva* sp. offshore the EMS with an onshore nutrient supply and developed a better understanding of seasonal growth dynamics and environmental effects (nitrogen, waves, depth, etc.). Specifically, we found that *Ulva* sp. is better cultivated deeper due to surface waves (i.e., 5m depth). In the EMS, sufficient nutrients should be provided before offshore cultivation. We developed a predictive model and validated it with data from our offshore experiments. Although in its infancy, this model has the potential to be used throughout the whole lifecycle of seaweed cultivation: from early nursery and farm design through economic models and estimations of environmental effects required by authorities and industry, ending with optimization of an ongoing farm operation. Future studies need to test the performance of other cultivation systems under similar conditions, examine alternative fertilization methods and validate the model on more offshore data from experimental and commercial farms. The model should be further improved by relating to P limitation, water motion, and waves.

The study allowed determining the impact of location, cultivation depth, nutrient availability, technology, and seasonality on the biomass yield and its chemical composition relevant for commodity production.

To progress the economically viable macroalgae-based supply chains, the follow-up study should translate the described technology into an analytical production function and investigate conditions for profitable production from a private and public perspective. The private perspective is based on market potential, while the public perspective considers the monetized external costs and benefits that might follow from macroalgae utilization.

## 5. Methods

### 5.1 Experimental part

#### 5.1.1 Marine macroalgae biomass

Starting at the autumn of 2018, *Ulva* sp. biomass maintained in Tel Aviv University system ^41^ was cultivated in a macroalgae photobioreactor (MPBR) built on a southern wall in the aquaculture center in Michmoret (Figure 5g). Nutrients were provided by constant exchange of nutrient rich seawater, pumped from the Michmoret bay. Throughout all experiments, the fresh weight (F.W.) of the biomass was determined using analytical scales after removing surface water using an electric centrifuge (Spin Dryer, CE-88, Beswin). Growth rates were calculated as Daily Growth Rate (DGR, eq. 1), as recommended by ^42^.

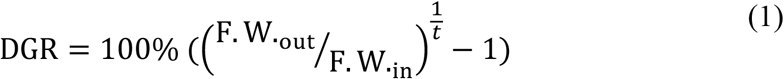

**Figure 5.**
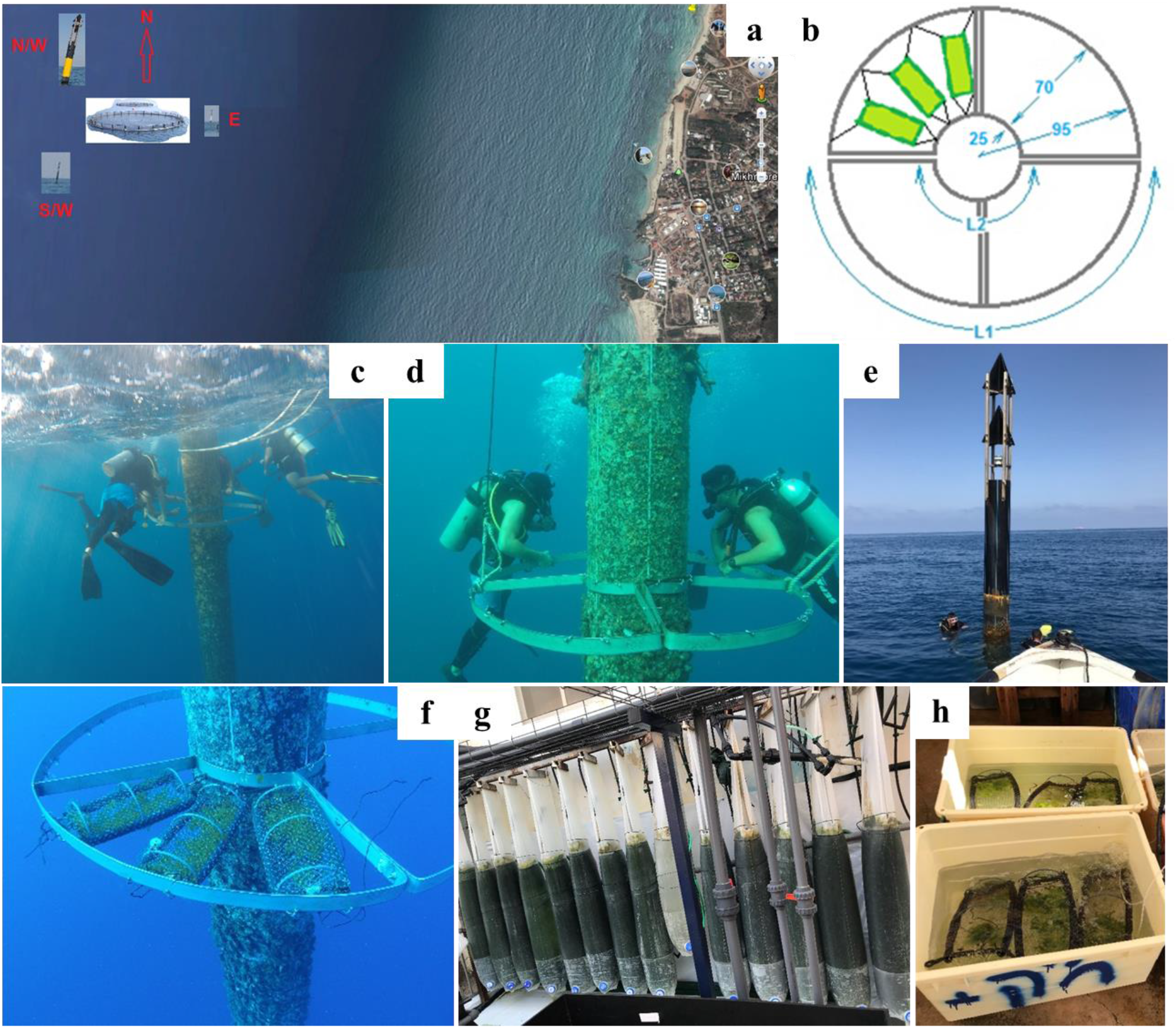
**(a)** Map of cultivation site offshore Michmoret, **(b)** illustration of a single cultivation ring and attached cages, **(c)** installation of the cultivation ring in a depth of 1m, **(d)** installation of the cultivation ring in a depth of 5m, **(e)** The North-Western marking buoy of the Lev-Yam fish cages, **(f)** three cages stocked with *Ulva* biomass installed on the cultivation system, **(g)** Image of the MPBR system used for maintaining *Ulva* sp. stocks in the aquaculture center in Michmoret, and **(h)** Onshore aerated fertilizing tanks, used between offshore cultivation periods.

Where FW_i_ (g) is the initial fresh weight, FW_out_ (g) is the final fresh weight and t is the number of cultivation days.

#### 5.1.2 Offshore cultivation systems

Cultivation experiments of *Ulva* sp. offshore Michmoret enabled us to measure year-round growth rates and chemical compositions in the Israeli EMS and to test the performance of a cultivation model for naturally varying environmental offshore conditions. The EMS offshore environment is ultra-oligotrophic, which also enabled assessing the effectiveness of short-term onshore fertilization as a method for recharging depleted nutrient storages in biomass between offshore cultivation periods.

Two ring-shaped offshore macroalgae cultivation systems (Figure 5c,d,f) were installed during November 2018 on a marking buoy of the Lev-Yam fish cages (Figure 5e) located 3.2 km offshore Michmoret (Figure 5a). The systems were welded from two flat 5mm*50mm stainless steel 316L rods bent into a large external ring (r = 0.95m), a small internal ring (r = 0.25m), and an internal connecting partial cross (Figure 5b). Each system was installed on the buoy by connecting both parts with stainless steel screws underwater. The systems were installed in two depths: 1m and 5m (Figure 5c-d). Each system was divided into four quarters and can carry up to twelve cultivation cages simultaneously (Figure 5b). The cages (0.15 m × 0.3 m, total illuminated area 0.045 m^2^) were built from polyethylene (D = 32 mm), high-density polyethylene (HDPE) (D = 16 mm) pipes, and two layers of polypropylene tubular nets (TENAX, Gallo Plastik, Italy) to allow full illumination and prevent grazing. The external nets were green 74N140 nets with a mesh size of 12-14. The internal nets were white 32G223 and 40G223 nets with mesh sizes of 15-20. They were fortified with a second layer of polyethylene net with smaller mesh holes to minimize biomass loss due to thalli fragmentation.

#### 5.1.3 Offshore cultivation experimental setup

Four successful experiments were performed between July 2019 and April 2020 (Table 3). A preliminary experiment was performed at the end of May 2019. In this experiment, the growth rates measured in the first cultivation period (after continuous fertilization) were invalid due to significant biomass losses due to large holes in the net. However, the growth rates measured in the second cultivation period were valid as we fortified the internal net between the first and the second cultivation periods.

**Table 3.** Details of offshore cultivation experiments.

|  | Cultivation<br>period 1* | Depth<br>[m] | Cages<br># | Depth<br>[m] | Cages<br># | Cultivation period<br>2* | Depth<br>[m] | Cages<br># | Depth<br>[m] | Cages<br># |
| --- | --- | --- | --- | --- | --- | --- | --- | --- | --- | --- |
| **Preliminary**<br>**experiment** | 20.5.19-28.5.19 |  | 1-6 |  | 13-18 | 29.5.19-6.6.19 |  | 1,3,4,<br>6 |  | 13-18 |
| **experiment 1** | 22-29.7.19 |  | 1-6 |  | 13-18 | 30.7.19-5.8.19 |  | 1-6 |  | 13-18 |
| **experiment 2** | 30.10.19-6.11.19 | 1 | 1-9 | 5 | 13-21 | - | 1 | - | 5 | - |
| **experiment 3** | 11.12.19-18.12.19 |  | - |  | 19-24 | 11.12.19-23.12.19** |  | - |  | 13-18 |
| **experiment 4** | 30.4.20-12.5.20 |  | - |  | 13-24 | - |  | - |  | - |

\*Cultivation period 1 began after a long-term continuous fertilization period, whereas cultivation period 2 began after a short one-day fertilization period.
\*\* In experiment 3, all cages were installed after continuous fertilization. The difference between period 1 and period 2 in this experiment is only cultivation duration.

At the beginning of each experiment, cages were stocked onshore with 20g F.W. of freshly harvested *Ulva* biomass from the Michmoret MPBR (Figure 5g), stitched tightly, transported by boat to the site and installed on the system by scuba divers (Figure 5e). The time that the biomass was outside water was kept to an unavoidable minimum. On the harvesting day, the cages were collected by scuba divers and returned to shore, where all biomass was taken out of the cages manually for weighing and storage for further chemical analysis. Finally, cages were cleaned thoroughly and stored for the next experiment.

As described in Table 3, different experiments put an emphasis on different aspects of offshore cultivation. In the preliminary and the first experiments, we examined whether a day of fertilization after a week of cultivation offshore the EMS (starvation conditions) supports further growth. This was investigated by performing two consecutive offshore cultivation periods separated by 18-24 hours of onshore fertilization. In the third experiment, performed during the winter, we examined if longer cultivation periods could be applied without compromising the daily yield. In this experiment, six cages were collected after seven days and six were collected after 12 days. The rationale of this design was that during the winter, there are fewer daily hours of light, and more extended cultivation periods may be needed to fulfill the biomass production potential.

#### 5.1.4 Onshore fertilization

A dedicated onshore experiment examined the effectiveness of 24-hours of fertilization of low nitrogen *Ulva* biomass from EMS offshore cultivation. The experiment tested if soaking low-N *Ulva* biomass after a week offshore, in a high-nutrient solution for 18-24 hours can recharge enough N to enable an additional week of growth. A separate experiment was performed to follow the increase in internal N during a 24 hours fertilization activity of starved *Ulva*. This experiment did not lead to interpretable results and is described in Appendix A.

The fertilization experiment was performed in two repetitions, during the preliminary cultivation experiment and experiment 1. Fertilization was applied in outdoor aerated 35L tanks, filled with nutrient-rich seawater. Half of the tanks were enriched in additional 1000 μM NH_4_ and 200 μM PO_4_ (Figure 5h). Cages were collected from the offshore system on the morning (9:00/11:00 of the 28.5.2019/29.7.2019). Fertilizing started at 13:00/16:00 after weighing, harvesting and restocking, and continued for 18-24 hours. The weight of the restocked biomass varied according to the amount of losses during the first cultivation period. Cages were divided into two fertilization groups as described in Table S3. After the fertilization period, the cages were returned offshore for an additional cultivation period. Water was sampled in the fertilization tanks at the end of each fertilization period. Finally, the fertilization effectiveness was inferred from the growth rate and chemical composition measured in the consecutive offshore cultivation period.

#### 5.1.5 Water sampling

Water sampling was performed in plastic syringes, filtered through a 0.2 μm filter to prevent particulate and microbial contamination, and kept at -20°C until analysis. Offshore water was sampled in every installation or collection of cages, at a 5m depth (by scuba divers) and at the sea surface (from the boat), representing 1m depth. Water was also sampled during the fertilization experiments, as described above.

#### 5.1.6 Temperature and irradiance

Temperature and irradiance were measured in each offshore experiment by a Onset® HOBO® sensor UA-002-08 (Onset Inc. MA), placed in one representative cage in each depth (Figure S2). The device also collected data during the onshore fertilizing period. In addition, as a backup for cases in which the HOBO devices were damaged or lost in the sea (Table S4), we used two more sources: 1. irradiance data from all experiment periods was extracted from the IMS data base from the Israel Meteorological Services (https://ims.data.gov.il/ims/1), and 2. water temperature was extracted from the Israel Marine Data Center (ISRAMAR) station located at a depth of 12m, 2.3 km offshore Hadera and 8 km North of the cultivation site in Michmoret. The suitability of the ISRAMAR data for the offshore model was determined based on a comparison to the available temperature measurements from the HOBO devices.

#### 5.1.7 Biomass chemical composition analysis

At the end of each cultivation experiment, biomass samples were harvested, weighed (F.W.), dried in 40-60°C, grinded with a mortar and pestle (and using liquid nitrogen if needed), and then kept at 4°C until further analysis. ∼70% of the samples underwent elemental analysis.

#### 5.1.8 Elemental Analysis

Elemental analysis for C, H, N, and S content as % of D.W. was performed at the Technion, Chemical, and Surface Analysis Laboratory, using Thermo Scientific CHNS Analyzer (Flash2000).

#### 5.1.9 Water analysis (Ammonia determination)

Ammonia concentration in water samples was determined following the method of Holms *et* al. (1999)^43^. Water samples were diluted using ultra-pure water aiming for concentrations lower than 0.5 μM NH_4_, which are optimal for this method.

#### 5.1.10 Data analysis

##### Fertilizing treatment

The effect of continuous and rapid fertilization on growth rates and internal N were compared using the two-tailed Mann-Whitney U test (groups # = 2, DF = 1).

##### Cultivation depth

The effect of cultivation depth, specifically 1m and 5m, on growth rates and internal N, were compared using the two-tailed Mann-Whitney U test (groups # = 2, DF = 1).

##### Cultivation Date

Growth rates and internal N in different dates were compared using the Kruskal-Wallis H test (groups # > 2, DF > 1), followed by the post-hoc Dunn’s test with the Bonferroni adjustment method for pairwise comparison.

##### Cultivation duration

The effect of cultivation duration, specifically 7 and 12 days, on growth rates and internal N, was assessed using the two-tailed Mann-Whitney U test (groups # = 2, DF = 1).

##### Correlation between DGR and internal N

A two-tailed Pearson test was used to compare and test the correlation between DGR and internal N results.

##### Analysis tools

Statistical analysis was performed using Python (3.7.3), specifically the scipy folder (1.4.1).

### 5.2 Model

Our model is based on the *Ulva* sp. dynamic cultivation model developed by ^39^. The model focuses on reactor scale *Ulva* sp. cultivation in offshore conditions and was constructed to study environmental effects on internal N and biomass growth dynamics in the offshore environment.

#### 5.2.1 Model Assumptions

The model follows the basic assumptions of the original model, developed by Zollmann et al. (2021) in ^39^, specifically that the dynamics of biomass growth and chemical composition are predicated by the dynamics of the limiting nutrient, in the case of the EMS nitrogen (N), under the constraining effect of light intensity (*I*). In the modeled environment, light intensity and temperature are varying in a daily and seasonal manner, but salinity is relatively constant and was assumed to be 39 PSU. Therefore, the salinity (S) function was removed from the model (fS = constant), as constant S has no effect on the system’s dynamics.

Following the concepts of the Droop Equation^44^, we assume that the effect of the concentration of the external N in the sea (*N*_*ext*_) on growth rate is not direct, but is mediated by the internal N in the biomass^22,45^. On the other direction, we assume that the biomass does not affect *N*_*ext*_, an assumption which will need to be reexamined in larger scales.

Our model also assumes that the organic carbon reserve, accumulated during the photosynthesis process, is not limiting within the modelled conditions and that all growth occurs during the light-period.

#### 5.2.2 Model Governoong Equations

The model is based on two governing ordinary differential equations (ODEs), describing the mass balance of two state variables: biomass density in the cage (m, g Dry Weight (D.W.) L^-1^, eq. 2) and biomass internal concentration of N (*N*_int,_ % gN gDW^−1^, eq 3), under constant *N*_*ext*_ and salinity. Both ODEs were solved numerically with hourly time steps.

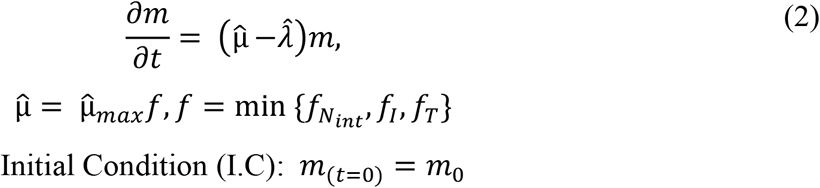

Where 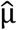 is the growth rate function in the offshore cage, 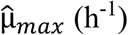 is the maximum specific growth rate under the applied salinity (39 PSU) conditions, and *f* is the combined growth function, made of 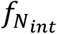 (eq 2.1) and *f*_*I*_ (eq 2.2), which are the *N*_int,_ and *I* growth functions. 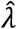 (eq. 2.1) is the biomass specific losses rate as at S= 39*PSU*. 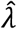 does not relate to losses in sporulation events. As described in ^39^, all rates appear on a per hour basis.

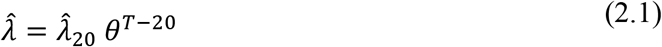

Where 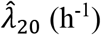 (h^-1^) is the specific rate of biomass losses and *θ* is an empiric factor of biomass losses.

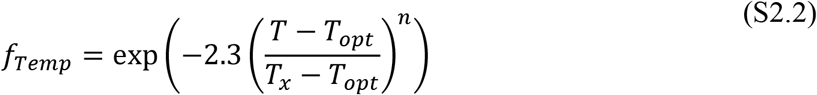

Where *T*_*x*_ = *T*_*min*_ for *T* ≤ *T*_*opt*_ and *T*_*x*_ = *T*_*max*_ for *T* > *T*_*opt*_. *T*_*min*_, *T*_*opt*_ and *T*_*max*_ (°C) are the minimal, optimal and maximal temperatures for *Ulva* growth.

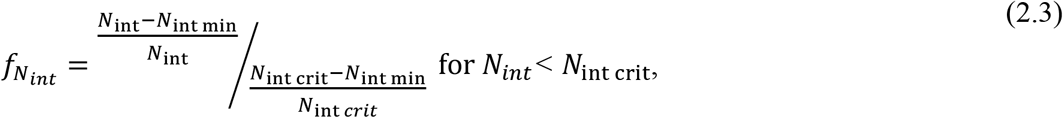

or 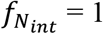 for *N*_*int*_ > *N*_int crit_

Where *N*_int min_ and *N*_int ma*x*_ (% g N g DW^-1^) are the minimum and maximum internal N concentrations in *Ulva*, respectively, *N*_*crit*_ (% g N g DW^-1^) is the threshold *N*_int_ level below which the growth rate slows down.

*f*_*I*_ (eq 2.2) was adjusted from the original light-function (eq. 6 in ^39^) as: 1. The z dimension of a flat cage is very small, and 2. the system was located 5 meters under the sea surface, requiring an additional light absorption term. Therefore, we added light extinction in the water column and assumed that light extinction in the water inside the cage is negligible.

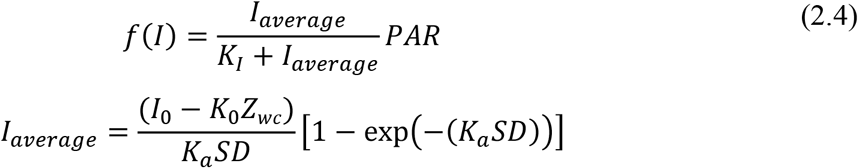

Where *I*_*average*_ and *I*_0_ (μmol photons m^−2^ s^−1^) are average photon irradiance in the cage and incident photon irradiance at water surface, respectively, *SD* (g D.W. m^-2^) is stocking density of biomass per unit of water surface in the cage, *K*_0_ (m^-1^) is the light extinction coefficient in the water, *Z*_*wc*_ (m) is water column depth above the cage and *K*_*a*_ (m^2^ g^-1^ DW) is the *Ulva* light extinction coefficient.

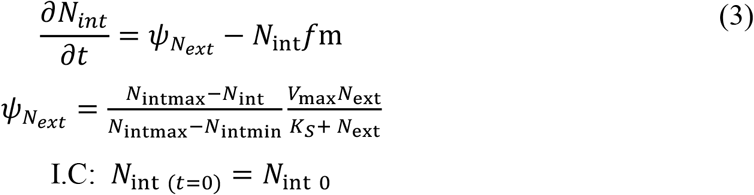

Where 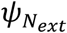 (μmol-N gDW h) is the N uptake function, formulated of *N*_intmax_ and *N*_intmi*n*_ (% gN gDW^−1^), *V*_m*ax*_ (μmol-N gDW^-1^ h^-1^), the maximum N uptake rate and *K*_S_ (μmol-N l^-1^), the N half-saturation uptake constant. −*N*_int_*f*m describes *N*_*int*_ dilution in biomass by growth.

#### 5.2.3 Model Parameters

Model parameters (Table S5) were determined in a three-steps calibration process, using a Sequential Least SQuares Programming optimizer (scipy) to minimize the Root Mean Square Relative Error (RMSRE) between model predictions and experimental data. First, parameters 1-10 were determined based on data from experiments of *Ulva* cultivation under various fertilizing regimes in a photobioreactor with controlled light and constant temperature and salinity^46^. Second, parameters 11-16 were determined based on data from experiments of *Ulva* cultivation in various mixes of nitrate-rich desalination brine and Artificial Seawater (ASW) (Appendix E). It should be noted that steps 1 and 2 of the calibration process was performed using the model with its original light function (eq. 6 in ^39^), before it was adjusted to the offshore system. Third, parameter 17, *K*_a_, and the *N*_ext_ levels during each experiment were evaluated by minimizing the average RMSRE between *m* and *N*_int_ for each experiment (*N*_ext_) and for all experiments together (*K*_a_).

#### 5.2.4 Model Simulations

Due to a lack of data regarding the exact levels of *N*_*ext*_, we estimated *N*_*ext*_ for each cultivation experiment by running the model for multiple *N*_*ext*_ levels and choosing the value with the smallest RMSRE between predictions and measurements. In some cases, in which predictions and measurements did not converge, we had to split the modeled cultivation period to a few shorter periods with different *N*_*ext*_ levels. This was done while considering the changing current regime that could potentially lead to nutrient enrichment in specific days of the cultivation period. Finally, we simulated biomass and *N*_*int*_ dynamics for each experiment, presenting predicted and measured levels of biomass and *N*_*int*_ at the end of each experiment.

## Acknowledgments

This work was supported by the Research Grant Award No. *US-5231-20CR* from BARD, The United States - Israel Binational Agricultural Research and Development Fund and by the Israel Ministry of Health (award # 3-16052). M.Z. thanks the Israeli Water Authority for partial funding of this study. The authors thank Rafi Yavetz, Carmel Gat and the students from the aquaculture center in the Ramot-Yam high school for accommodating the study and providing offshore and underwater logistical support, prof. Gitai Yahel from the Oceanography and Marine Biology Laboratory in the Ruppin academic center for assisting with ammonia measurement, and Mr. Alex Chemodanov for designing and constructing the offshore system.

