## Supplementary material for "Cultivation of *Ulva* sp. offshore the Eastern Mediterranean Sea in experimental bioreactors: seasonal growth dynamics and environmental effects"

### Appendix A - Onshore fertilizing in bottles

#### Introduction

The goal of this Appendix is to describe an onshore fertilizing experiment that was performed after a period of offshore *Ulva* sp. cultivation, to follow the increase in its internal N during a 24-hours fertilization activity. This experiment was performed to collect complementary data on the effectiveness of rapid onshore fertilizing between two periods of offshore cultivation in the oligotrophic EMS. However, as the results were not interpretable, this part was excluded from the main text.

#### Methods

This 24-hours fertilization experiment was performed after experiment #2 cultivation period, starting during the 6.11.2019. fertilization was performed in outdoor aerated 1.5L bottles, filled with seawater and enriched in additional 500 µM $NH_4$ and 50 µM P (Figure S1). Half of the bottles were shaded by a 50% cover net, and half were not shaded. Cages were collected from the offshore system on the morning (11:00) of the 6.11.2019. Fertilization started at 16:00 after harvesting and weighing. Biomass was taken from two cages, one from each cultivation depth and was fertilized in shaded or unshaded bottles in a stocking density of 1 g FW $L^{-1}$ in triplicates, as describe in Table S1. Water was sampled during the following day at 9:00, 12:00 and 15:00 for ammonia analysis. Small biomass pieces (~0.2g F.W.) were sampled for CHNS analysis at the same time points.

**Table S1. Details of second fertilization experiment**

| Cage # | Cultivation depth [m] | Bottle # | Light/shade |
|---|---|---|---|
| 1 | 1 | a | Light |
| | | b | Shade |
| | | c | Light |
| | | d | Shade |
| | | e | Light |
| | | f | Shade |
| 13 | 5 | a | Light |
| | | b | Shade |
| | | c | Light |
| | | d | Shade |
| | | e | Light |
| | | f | Shade |

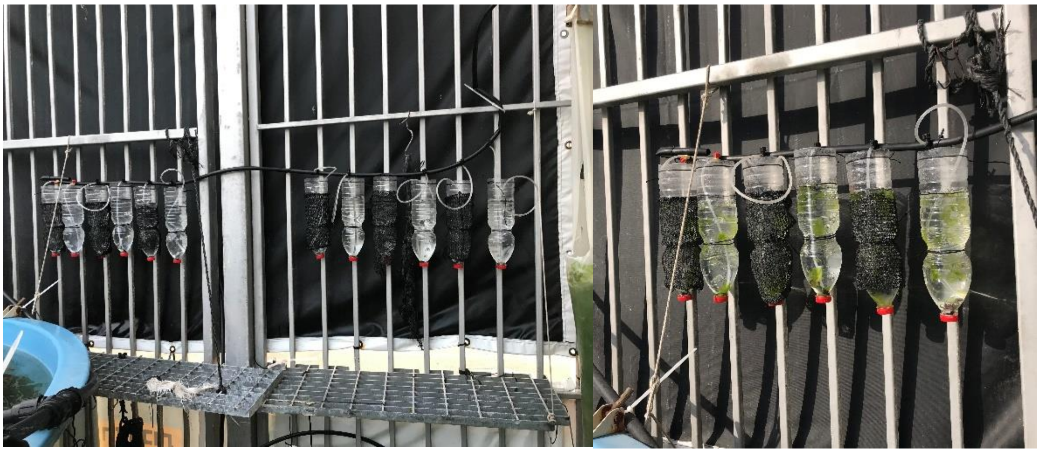

**Figure S1.** Onshore bottle fertilization

### Results and discussion

Levels of ammonium in water and of internal N in biomass samples were quantified at 9:00, 12:00 and 15:00 of the 7.11.2019, which are 17, 20 and 23 hours since fertilizing started. All measurements are available in Table S2. However, as the results of ammonia quantification were not deemed fit for evaluation, we cannot conclude anything quantitative from these results beyond the clear decreasing trend. The internal N measurements, on the contrary, are supposed to be reliable, but the sample size and the trends somewhat contradicting figures (i.e., a decrease in internal N during a fertilizing process) do not allow us to offer any interpretation.

Beyond the unclear results, the experimental design is weak as the biomass sampling process, necessary for internal N quantification, effects significantly the biomass stocking density and poses a potentially significant external effect on the fertilizing process.

In conclusion, the timewise effect of level of shading on uptake rate and recovery of internal N should be examined in a better designed experiment, with a larger sample size and larger cultivation systems. Otherwise, small cultivation systems can be used to examine initial and final internal N levels, without sampling along the experiment.

**Table S2. Results of onshore bottles fertilizing**

| Offshore cultivation depth [m] | bottle # | Shade / light | Measured ammonium (µM $NH_4$) | | | Measured internal N (% g N $g^{-1}$ DW) | | |
|---|---|---|---|---|---|---|---|---|
| | | | 17 hours | 20 hours | 23 hours | 17 hours | 20 hours | 23 hours |
| 1 | a | light | 382 | 401 | 310 | 1.65 | 2.16 | 1.77 |
| | b | shade | 752 | 428 | 301 | 1.63 | 1.70 | |
| | c | light | 485 | | 442 | | | |
| | d | shade | 668 | | 350 | | | |
| | e | light | 347 | | 246 | | | |
| | f | shade | 485 | | 346 | | | |
| 5 | a | light | 447 | 408 | 382 | 1.63 | 1.70 | 1.92 |
| | b | shade | 409 | 287 | 292 | 1.77 | 2.44 | 2.49 |
| | c | light | 399 | | 216 | | | |
| | d | shade | 517 | | 351 | | | |
| | e | light | 396 | | 240 | | | |
| | f | shade | 401 | | 228 | | | |

### Appendix B - Details of onshore tank fertilizing

In the preliminary experiment and in experiment #1 the rapid onshore fertilization between the two cultivation periods included two groups. One group of cages was fertilized with seawater with no artificial fertilizer addition and one group was fertilized in seawater supplemented with 1000 µM $NH_4$ and 200 µM $PO_4$. The full details are described in Table S3. At the end of the experiments, no difference was found between those two groups regarding the daily growth rate (DGR).

**Table S3. Details of tank fertilization experiment**

| Experiment # | Fertilization duration | Fertilization medium | Cage # | Cultivation depth [m] | Weight of fertilized biomass [g F.W.] |
|---|---|---|---|---|---|
| Preliminary | 24 hours (13:00 28.5.19- 13:00 29.5.19) | Seawater | 4 | 1 | 1.2 |
| | | | 6 | 1 | 2.7 |
| | | | 16 | 5 | 6.6 |
| | | | 17 | 5 | 6.6 |
| | | | 18 | 5 | 11.9 |
| | | Seawater + 1000 µM NH4 + 200 µM $PO_4$ | 1 | 1 | 4.4 |
| | | | 3 | 1 | 4 |
| | | | 13 | 5 | 8.4 |
| | | | 14 | 5 | 6.5 |
| | | | 15 | 5 | 5.5 |
| 1 | 18 hours (16:00 29.7.19- 10:00 30.7.19) | Seawater | 4 | 1 | 16 |
| | | | 5 | 1 | 20 |
| | | | 6 | 1 | 20 |
| | | | 16 | 5 | 20 |
| | | | 17 | 5 | 19.9 |
| | | | 18 | 5 | 20 |
| | | Seawater + 1000 µM NH4 + 200 µM $PO_4$ | 1 | 1 | 16 |
| | | | 2 | 1 | 15.8 |
| | | | 3 | 1 | 19.8 |
| | | | 13 | 5 | 19.9 |
| | | | 14 | 5 | 20.1 |
| | | | 15 | 5 | 20.2 |

### Appendix C - HOBO devices in cages

Figure S2 presents the HOBO device used to measure light and temperature during the offshore experiments, inside and outside the cultivation cage. Table S4 shows the details of HOBO devices placement and fate (if damaged) in the different experiments.

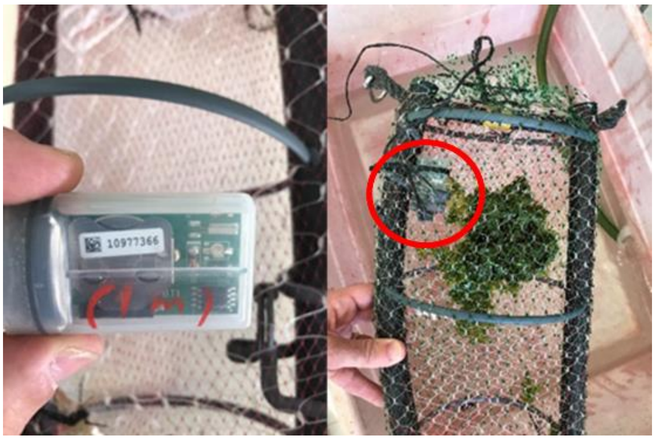


**Figure S2**. HOBO device for measurement of light and temperature, outside **(Left)** and inside **(Right)** a cultivation cage

**Table S4. HOBO devices placement in the different experiments**

| Run | Depth [m] | Cage # | Damaged? | Depth [m] | Cage # | Damaged? |
|---|---|---|---|---|---|---|
| Preliminary - first period | 1 | - | - | 5 | - | - |
| Preliminary - second period | 1 | 1 | No | 5 | 13 | Yes |
| #1 (both periods) | 1 | 1 | No | 5 | 13 | No |
| #2 | 1 | 1 | No | 5 | 13 | No |
| #3 | - | - | - | 5 | 13 | Yes |
| #4 | - | - | - | 5 | - | |

### Appendix D - Model parameters

Table S5 presents the details of all model parameters, which were determined in a three-steps calibration process as described in the model chapter in the main text.

**Table S5. Model parameters**

| Parameter # | Parameter symbol | Calibration system | Value | Unit |
|---|---|---|---|---|
| 1 | $\hat{\mu}_{max}$ | Indoor controlled photobioreactor[46] | 0.03 | Light $h^{-1}$ |
| 2 | $T_{min}$ | | 4 | ℃ |
| 3 | $\hat{\lambda}$ | | 0.003 | Light $h^{-1}$ |
| 4 | $N_{int\ min}$ | | 0.48 | % g N $g^{-1}$ DW |
| 5 | $N_{int\ max}$ | | 4.5 | % g N $g^{-1}$ DW |
| 6 | $N_{int\ crit}$ | | 0.7 | % g N $g^{-1}$ DW |
| 7 | $K_s$ | | 10 | μmol N $L^{-1}$ |
| 8 | $V_{max}$ | | 50 | μmol N $g^{-1}$ DW $h^{-1}$ |
| 9 | $K_I$ | | 15 | μmol photons $m^{-2}$ $s^{-1}$ |
| 10 | $K_0$ | | 0.2 | $m^{-1}$ |
| 11 | $S_{min}$ | outdoor semi-controlled bottles photobioreactor (Appendix C) | 0 | PSU |
| 12 | $S_{opt}$ | | 28 | PSU |
| 13 | $S_{max}$ | | 50 | PSU |
| 14 | $T_{opt}$ | | 18 | ℃ |
| 15 | $T_{max}$ | | 36 | ℃ |
| 16 | n | | 5.1 | − |
| 17 | $K_a$ | Offshore system | 0.2 | $m^2$ $g^{-1}$ DW |

### Appendix E - outdoor semi-controlled bottles photobioreactor

### Introduction

The outdoor semi-controlled bottles photobioreactor was established as part of study aimed to examine the effectiveness of *Ulva* sp. cultivation in nitrate rich ground water desalination brine for the sake of nitrate removal and biomass production. This study included an experimental part, in which *Ulva* was cultivated in various dilutions of brine and ASW and different stocking densities, measuring growth rates and internal N content at the beginning and the end of each experiment, and a modeling part, in which the reactor scale cultivation model was adjusted to this system.

### Methods

### Cultivation system

Twelve upside down 1.5L transparent PET bottles, positioned on the Southern wall of the Porter building in TAU, were used as photobioreactors (Figure S3). Each reactor was filled up with 1 L composed of a dilution of ASW (39 PSU) and nitrate rich ground water desalination brine (6 PSU, 265 mg- $NO_3^-$ $L^{-1}$), provided from the Israeli water company, Mekorot, from the Ashkelon brine facility. The bottles were well mixed by bottom aeration and no water exchange was performed.

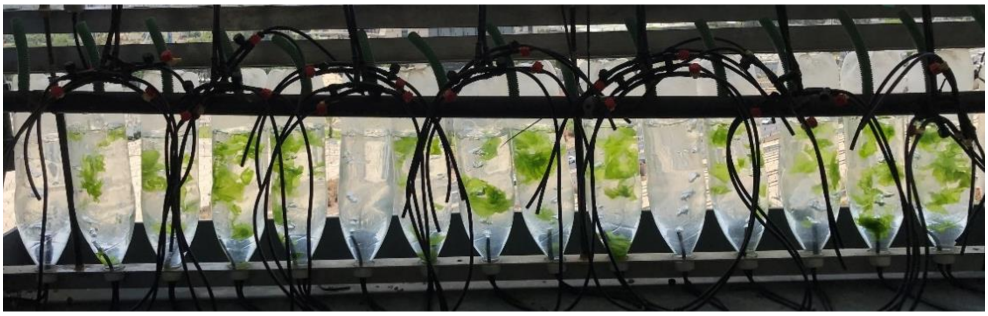

**Figure S3**. Outdoor bottles photobioreactors stocked with Ulva thalli during a cultivation experiment in various dilutions of nitrate rich ground water desalination brine and ASW.

### Experimental setup

*Ulva* sp. was cultivated in the outdoor bottles photobioreactors for two consecutive periods of 6-7 days during June 2020, examining the effects of four dilution ratios of ASW and nitrate rich brine and three stocking densities (1, 2 and 3 g F.W. $l^{-1}$), as described in Table S6. Phosphorus was not supplemented to the cultivation reactors but is expected to be required for longer cultivation periods.

Experiment measurements focused on growth rate and N content (CHNS analysis), while the environmental parameters temperature and irradiance were monitored for control by a HOBO device that was positioned in an adjacent bottle, filled only with ASW.

**Table S6. Experimental data used for calibration of outdoor semi-controlled bottles photobioreactor**

| experiment # | Bottle # | Dilution rate [L brine : L ASW] | Salinity [PSU] | Initial nitrate concentration [mg-$NO_3$ $L^{-1}$]* | Stocking density [g FW $L^{-1}$] | Final m [g FW $L^{-1}$] | Final Nint [% g N gDW] | Comment |
|---|---|---|---|---|---|---|---|---|
| 1** | 2,3 | 1:0 | 6 | 265 | 1 | 1.3, 2.1 | 3.1, 2.3 | |
| | 4,5 | | | | 2 | 4, 3 | 2.5, 2.8 | |
| | 7,8 | 0.5:0.5 | 24 | 132.5 | 1 | 3.8, 4.5 | 2.4, 2.1 | |
| | 9,10 | | | | 2 | 7.8, 8.1 | 1.6, 1.6 | |
| | 12,13 | 0.33:0.67 | 29 | 88.3 | 1 | 3.7 ,3.5 | 2, 1.9 | |
| | 14,15 | | | | 2 | 6.9, 5.3 | 1.4, 1.9 | |
| 2*** | 2,3 | 1:0 | 6 | 265 | 2 | 3, 2.6 | 2.9, 3.2 | |
| | 4,5 | | 40 | 265 | 2 | 5, 4.6 | 2.8, 2.6 | Salted brine |
| | 7,8 | 0.5:0.5 | 22.5 | 132.5 | 2 | 4.4, 5.7 | N/A, 2.2 | |
| | 9,10 | | | | 3 | 5.7, 5.8 | 2.9, 2.3 | |
| | 12,13 | 0.67:0.33 | 17.5 | 176.7 | 2 | 5.1, 4.7 | 2.9, 2.2 | |
| | 14,15 | | | | 3 | 6.6,6.9 | 2.8, 2.4 | |

*All concentrations are calculated based on the Mekorots report on a concentration of 265 mg $NO_3$ $L^{-1}$, and a negligible concentration in the ASW

** Nint,0 = 3.03 % g N $g^{-1}$ DW

** Nint,0 = 4.09 % g N $g^{-1}$ DW

### Appendix F – Growth rates and internal N data

Tables S7 and S8 summarize DGR and internal N results from the offshore experiments.

**Table S7. Daily growth rates measured for *Ulva* sp. in different offshore cultivation experiments**

| | Dates | Fertilizing regime* | Depth | N [# Samples] | Mean DGR [% day$^{-1}$] | S.D. [% day$^{-1}$] | S.E. [% day$^{-1}$] | 95% Conf. Interval [% day$^{-1}$] | |
|---|---|---|---|---|---|---|---|---|---|
| **Preliminary experiment** | 29.5.19-6.6.19 | Rapid | 1 | 4 | -25.6 | 11.3 | 5.7 | -43.6 | -7.6 |
| | | | 5 | 6 | -39.4 | 47.5 | 19.4 | -89.2 | 10.4 |
| **Experiment 1** | 22-29.7.19 | Continuous | 1 | 6 | 1.2 | 3.2 | 1.3 | -2.2 | 4.6 |
| | | | 5 | 6 | 9.8 | 2.4 | 1 | 7.2 | 12.4 |
| | 30.7.19-5.8.19 | Rapid | 1 | 6 | -19.3 | 11.1 | 4.6 | -31 | -7.6 |
| | | | 5 | 6 | -11.7 | 5.4 | 2.2 | -17.4 | -5.9 |
| **Experiment 2** | 30.10.19-6.11.19 | Continuous | 1 | 9 | -4.3 | 4.5 | 1.5 | -7.7 | -0.8 |
| | | | 5 | 9 | 6.1 | 0.8 | 0.3 | 5.4 | 6.7 |
| **Experiment 3** | 11-18/23.12.19 | Continuous | 5 | 12 | 0.6 | 3.1 | 0.9 | -1.3 | 2.6 |
| **Experiment 4** | 30.4.20-12.5.20 | Continuous | 5 | 12 | 7.3 | 2.4 | 0.7 | 5.8 | 8.8 |

*Fertilizing regime relates to nutrient enrichment prior to the cultivation experiment. Continuous fertilizing refers to prolong cultivation in the outdoor MPBR in Michmoret which receives nutrient rich near-shore water, whereas rapid fertilizing refers to a 18-24 hours of onshore fertilizing between two cultivation periods in the nutrient poor offshore environment.

**Table S8. Internal N measured for *Ulva* sp. in different experiments of the offshore cultivation experiment**

| | Dates | Fertilizing regime* | Depth | N [# Samples] | Mean internal N [% g N $g^{-1}$ DW] | S.D. [% g N $g^{-1}$ DW] | S.E. [% g N $g^{-1}$ DW] | 95% Conf. Interval [% g N $g^{-1}$ DW] | |
|---|---|---|---|---|---|---|---|---|---|
| **Preliminary experiment** | 20-28.5.19 | Continuous | 1 | 1 | 0.84 | - | - | - | - |
| | | | 5 | 2 | 0.63 | 0.16 | 0.11 | -0.78 | 2.03 |
| | 29.5.19-6.6.19 | Rapid | 1 | 1 | 0.83 | - | - | - | - |
| | | | 5 | 2 | 0.50 | 0.05 | 0.04 | 0.01 | 0.98 |
| **Experiment # 1** | 22-29.7.19 | Continuous | 1 | 2 | 0.91 | 0.27 | 0.19 | -1.52 | 3.34 |
| | | | 5 | 2 | 0.83 | 0.10 | 0.07 | -0.06 | 1.72 |
| | 30.7.19-5.8.19 | Rapid | 1 | 2 | 0.72 | 0.08 | 0.06 | -0.02 | 1.47 |
| | | | 5 | 2 | 0.68 | 0.11 | 0.08 | -0.30 | 1.67 |
| **Experiment # 2** | 30.10.19-6.11.19 | Continuous | 1 | 2 | 0.98 | 0.01 | 0.01 | 0.89 | 1.07 |
| | | | 5 | 2 | 1.05 | 0.18 | 0.13 | -0.57 | 2.66 |
| **Experiment # 3** | 11-18/23.12.19 | Continuous | 5 | 8 | 1.95 | 0.28 | 0.10 | 1.72 | 2.18 |
| **Experiment # 4** | 30.4.20-12.5.20 | Continuous | 5 | 5 | 1.64 | 0.13 | 0.06 | 1.48 | 1.81 |

*Fertilizing regime relates to nutrient enrichment prior to the cultivation experiment. Continuous fertilizing refers to prolong cultivation in the outdoor MPBR in Michmoret which receives nutrient rich near-shore water, whereas rapid fertilizing refers to a 18-24 hours of onshore fertilizing between two cultivation periods in the nutrient poor offshore environment.

### Appendix G - Daily growth rates and internal N after rapid and continuous nutrient enrichment in depths of 1 m and 5 m

Figures S4 compares the results of each depth (1 and 5 meters cages) separately (p-value < 0.001). This figure shows a consistent but relatively mild effect on internal N in both depths.

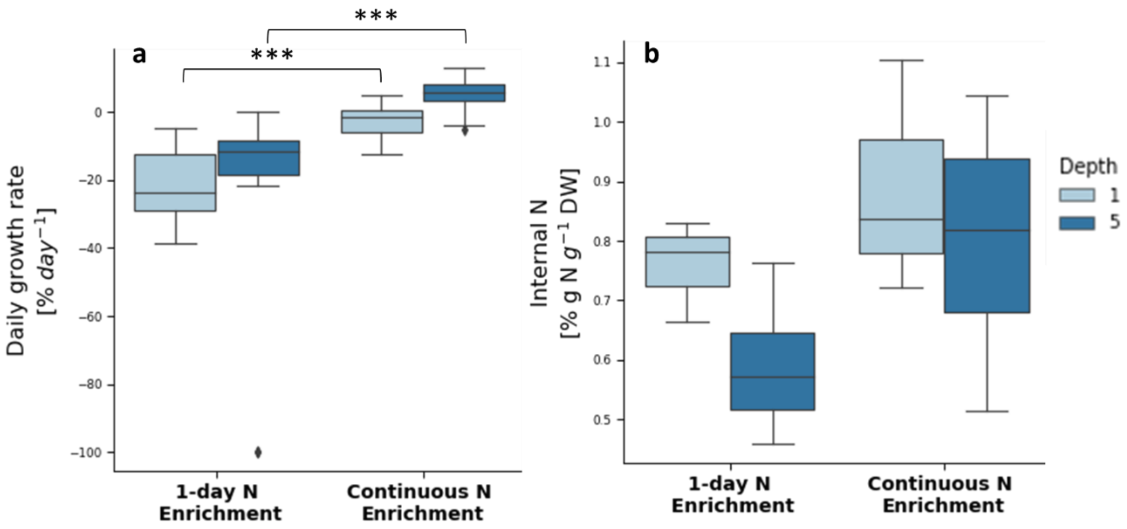


**Figure S4**. Daily growth rates **(a)** and internal N **(b)** of *Ulva* sp. cultivated offshore the EMS after rapid nutrient enrichment and after continuous nutrient enrichment in depths of 1 m **(light blue)** and 5 m **(dark blue)**, in the preliminary and in experiment #1**.** Asterisks indicate statistical significance of difference with *** $p < 0.001$, calculated by the two-tailed Mann-Whitney U test. # samples: rapid nutrient enrichment:  6 for DGR and 2 for internal N for each depth. continuous nutrient enrichment: 15 and 39 for DGR and 6 and 8 for internal N, for depth of 1 and 5 m, respectively.

### Appendix H – Images of the harvested cultivation cages

Figures S5 and S6 present images of the offshore cages after experiments #1 and #2, respectively, from cultivation in both depths (1 and 5 m). Figure S6 presents the images of all cages from experiments #1-4, cultivated in a 5m depth.

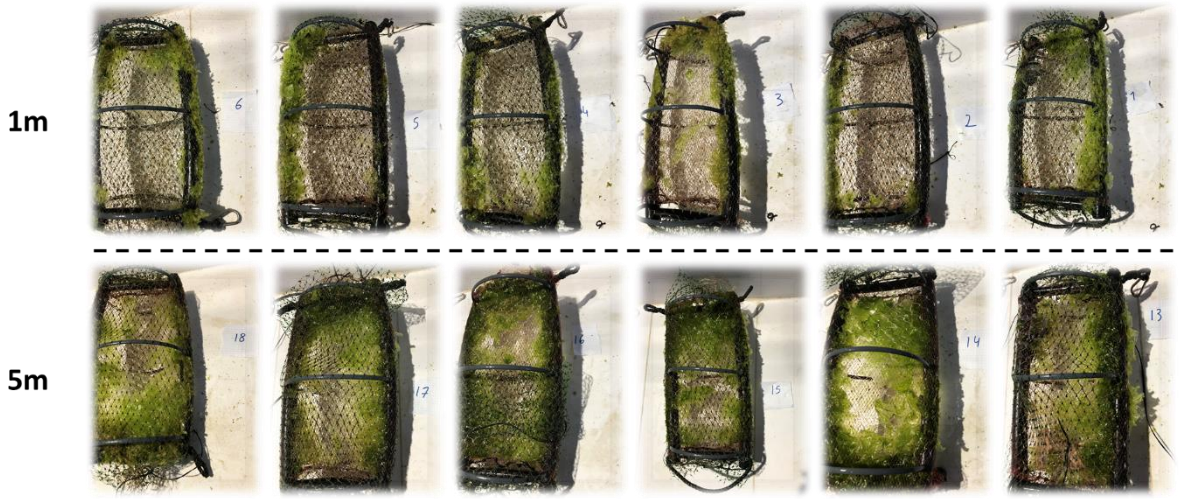


**Figure S5**. Offshore cages at the end of the cultivation period of experiment #1 (22-29 July 2019), in a depth of 1m **(top row)** and 5m **(bottom row).**

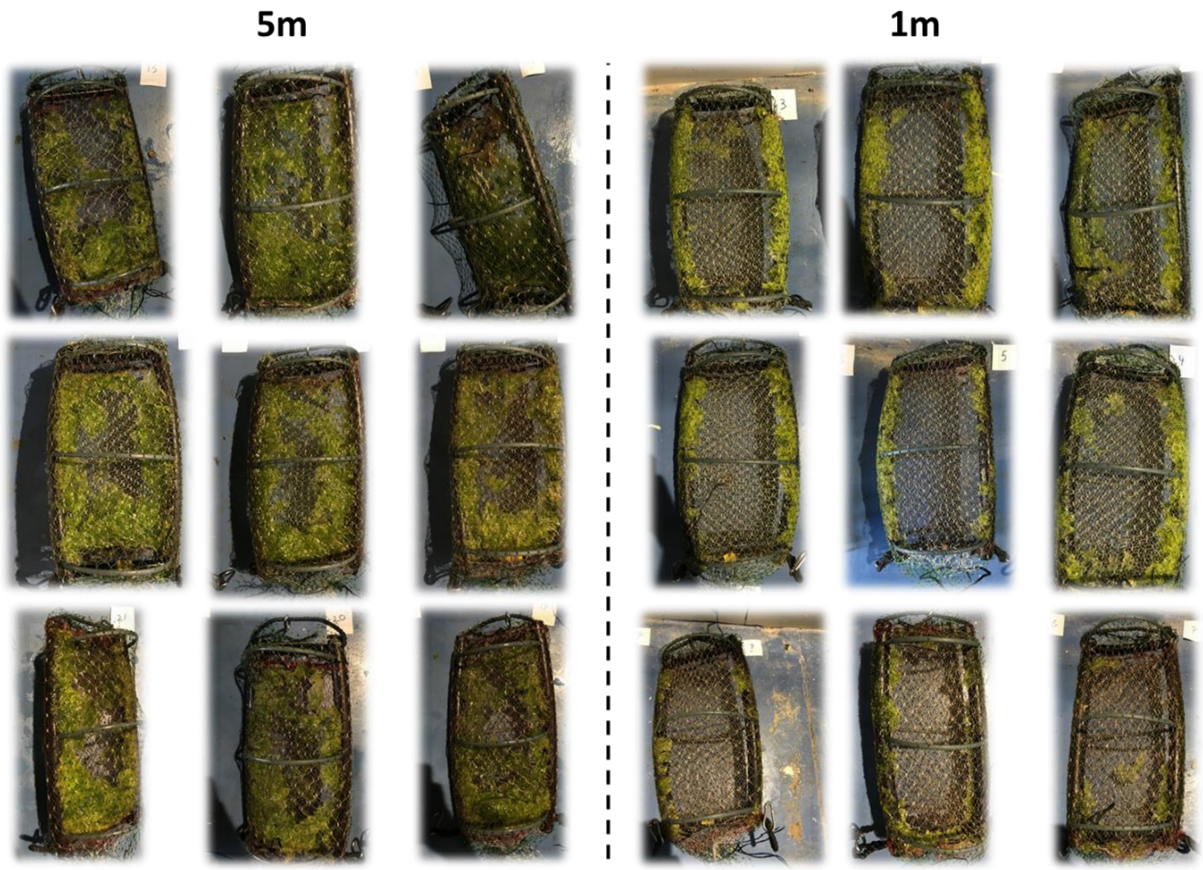


**Figure S6**. Offshore cages at the end of the cultivation period of experiment #2 (30 October – November 6 2019), in a depth of 1m **(right)** and 5m **(left).**

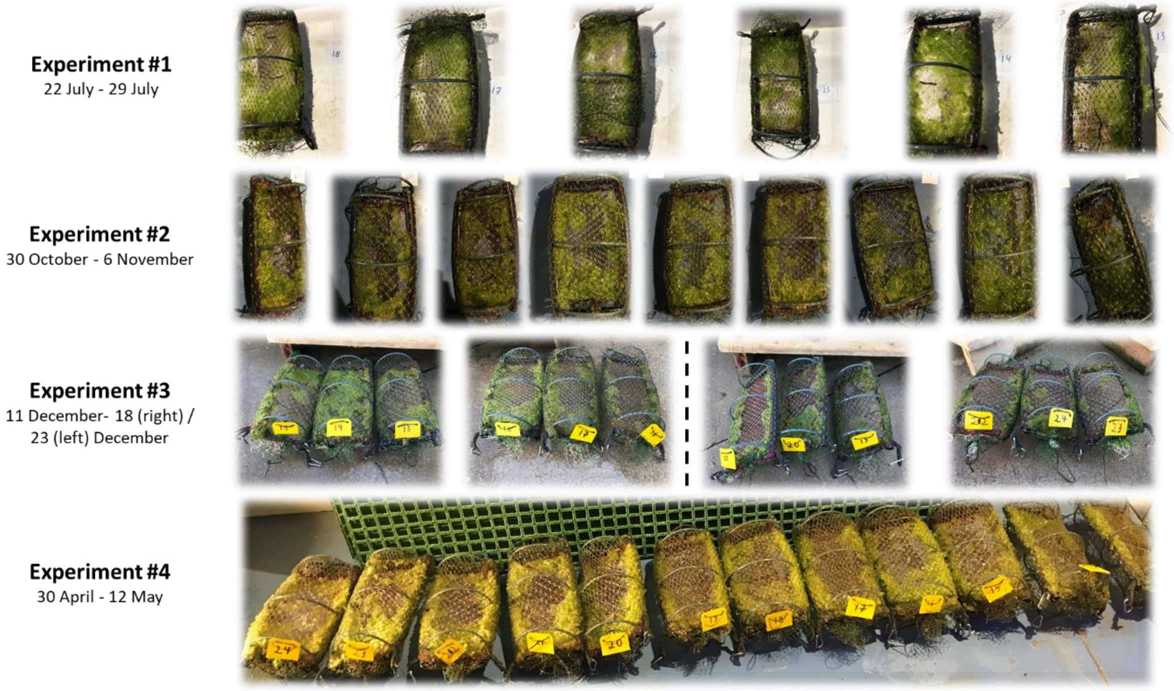


**Figure S7**. Offshore cages from 5m depth, after continuous fertilization, at the end of the cultivation periods. Each row presents cages from a different run, performed in a different season (from top down: summer, autumn, winter and spring). Cages of experiment #3 are divided between cultivation durations of 7 days (right) and 12 days (left).

### Appendix I - Offshore weather conditions during cultivation experiments

#### Introduction

*Ulva* cultivation offshore is highly dependent on meteorological conditions such as light intensity, water temperature, waves, currents, and winds. The goal of this Appendix is to provide graphic illustrations of those conditions, when relevant, to help and interpret the effect of those time dependent factors on results of the offshore *Ulva* sp. cultivation experiments.

#### Methods

This Appendix incorporates data from the IMS data base from the Israel Meteorological Services (https://ims.data.gov.il/he/ims/6) and from the IOLR ISRAMAR and Mediterranean GLOSS #80 station, located 2.3 km offshore Hadera. All data is plotted in per experiment plots.

#### Results

Light intensity and water temperature measurements along the offshore cultivation periods are presented among models simulations for experiments #1-2 and #4 (Figure 4) and herein in Figure S8. This data is used to interpret the results of the offshore cultivation experiments, and as an input to the offshore cultivation model.

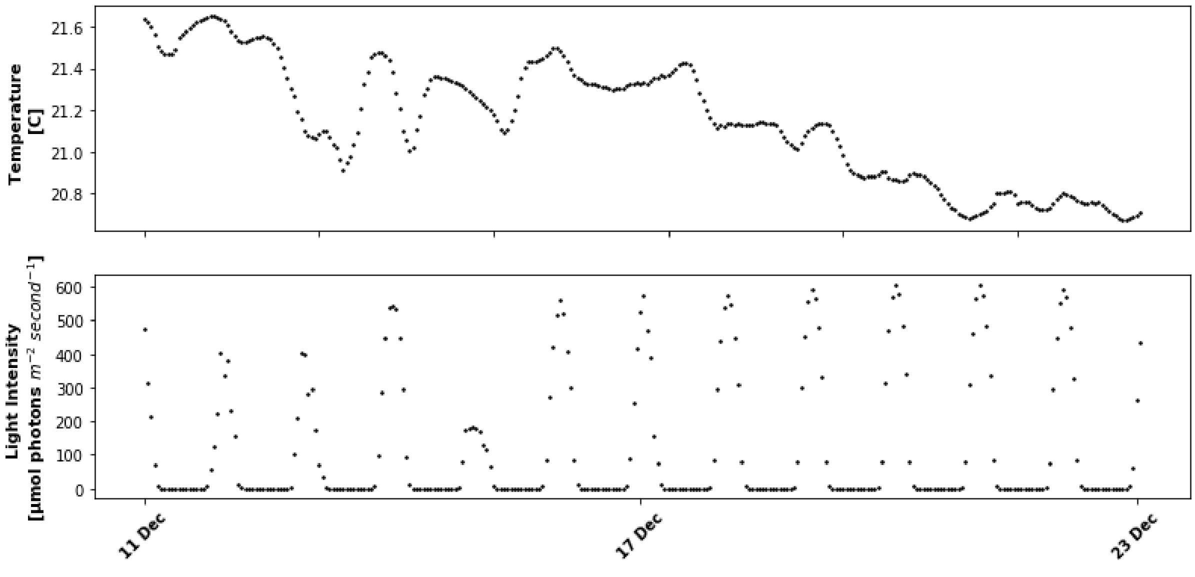


**Figure S8.** Water temperature **(top row)** and light intensity **(bottom row)** measured during experiment #3.

Significant wave heights (height of upper third of the waves) along the offshore cultivation periods are presented in Figure S9. Focusing on the periods we the higher waves (above 2 meters), we present in Figure S10 the wind directions during the days before the rising of the waves in experiments #3 (11.12.2019) and #4 (4-5.5.2020). The figures show that on the 11.12.2019, the dominant wind direction was easterly (western wind), whereas on the 4-5.5.2020 the dominant wind direction was westerly (eastern wind).

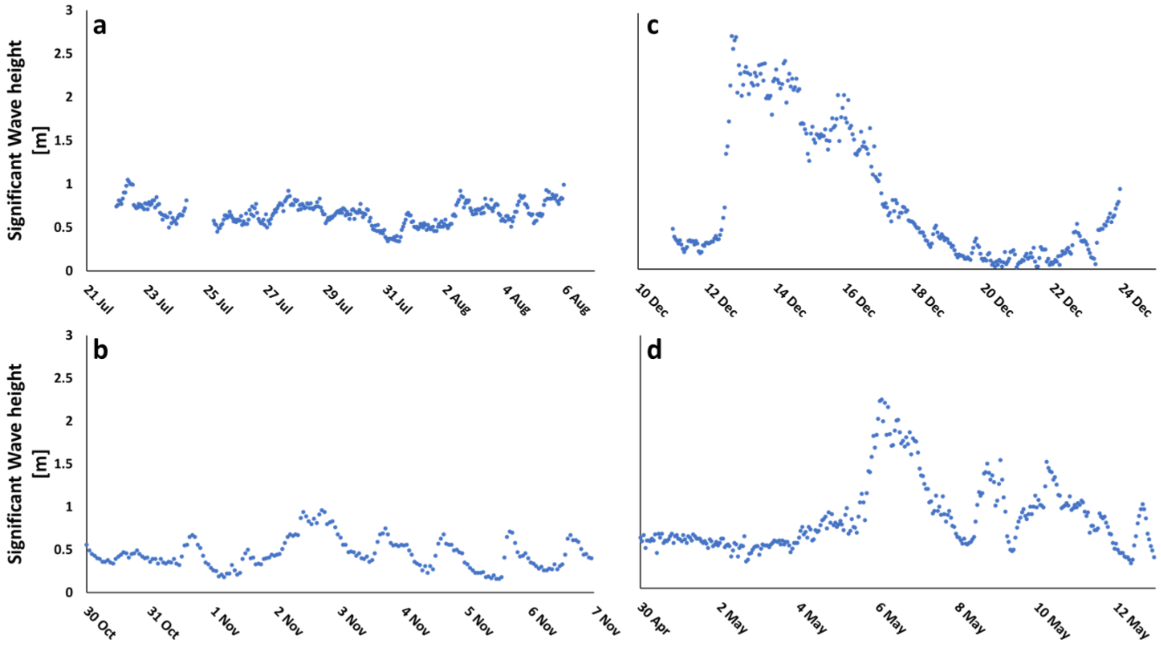


**Figure S9**. Significant wave heights during experiments #1 **(a)**, #2 **(b)**, #3 **(c)** and #4 **(d)** of the offshore *Ulva* sp. cultivation experiment, as measured in the Hadera GLOSS #80 station. Experiments #1-#3 were performed in 2019 and experiment #4 was performed in 2020.

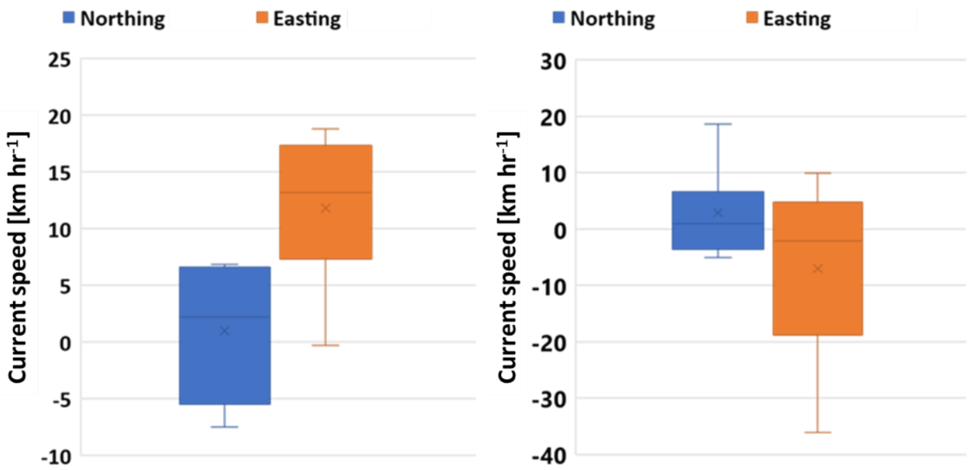


**Figure S10.** Wind direction and speed before the rising of the waves in experiment #3, during the 11.12.2019 **(left)**, and in experiment #4, during the 4-5.5.2020 **(right)**.

Current regime during the cultivation periods, potentially effecting N supply to the cultivation system by supplying nutrients from the fish cages or from the nutrient enriched Alexander estuary, is presented in Figure S11.

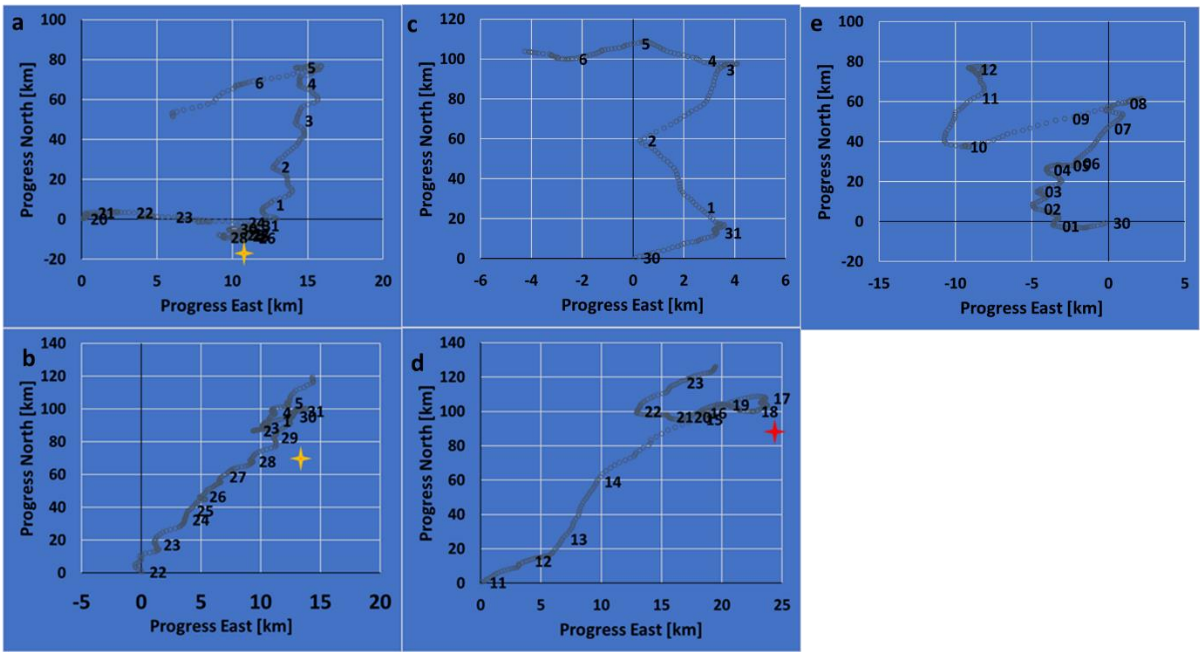


**Figure S11.** Current regime analysis of the preliminary experiment **(a)** and experiments #1 **(b)**, #2 **(c)**, #3 **(d)** and #4 **(e)** of the offshore *Ulva* sp. cultivation experiment, as measured in the Hadera GLOSS #80 station. Numbers on the plots represent the day of the month. Yellow stars in the preliminary experiment and in experiment #1 represent the harvesting day of the first cultivation period, after continuous fertilizing. The red star in experiment #3 represents the harvesting day of the first half of the cages in experiment #4.

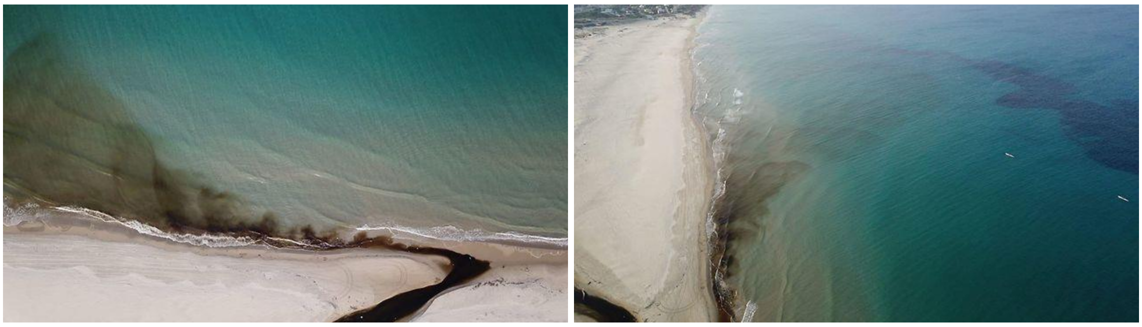

**Figure S12**. Aerial photo of the baseflow distribution of the the Alexander Estuary discharge during a heavy pollution event on the 6.11.2019 (photo credit: Erez Harari, YNET, https://www.ynet.co.il/articles/0,7340,L-5621116,00.html)

### Appendix J – Illustrated sensitivity of model results to model parameters

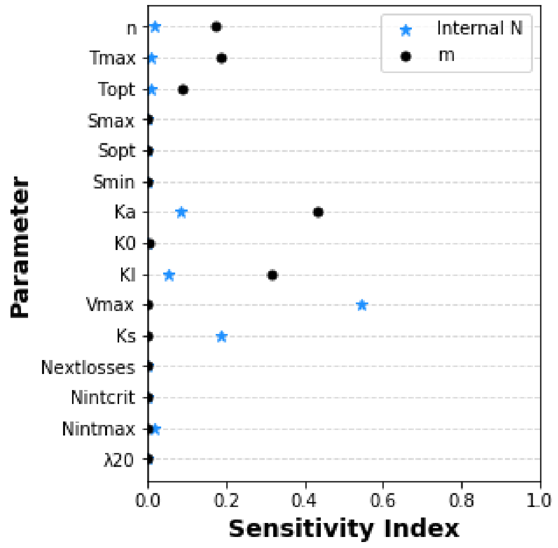


**Figure S13**. Illustrated sensitivity of simulated biomass production (black circles) and N content (blue stars) to model parameters, as measured by the Sobol method, in the offshore cultivation system**.**

### Appendix K – Model simulation of nitrogen and biomass dynamics under a short-term fertilizing treatment

Figure S14 presents models simulations of a high peak, short-term (20 hours) fertilization treatment in a closed system under controlled light, temperature, salinity and nutrient (N and P) enrichment. The simulation predicts the recovery of more than 1 % g N $g^{-1}$ D.W. and a significant growth in the following week, which were not observed in the offshore experiments.

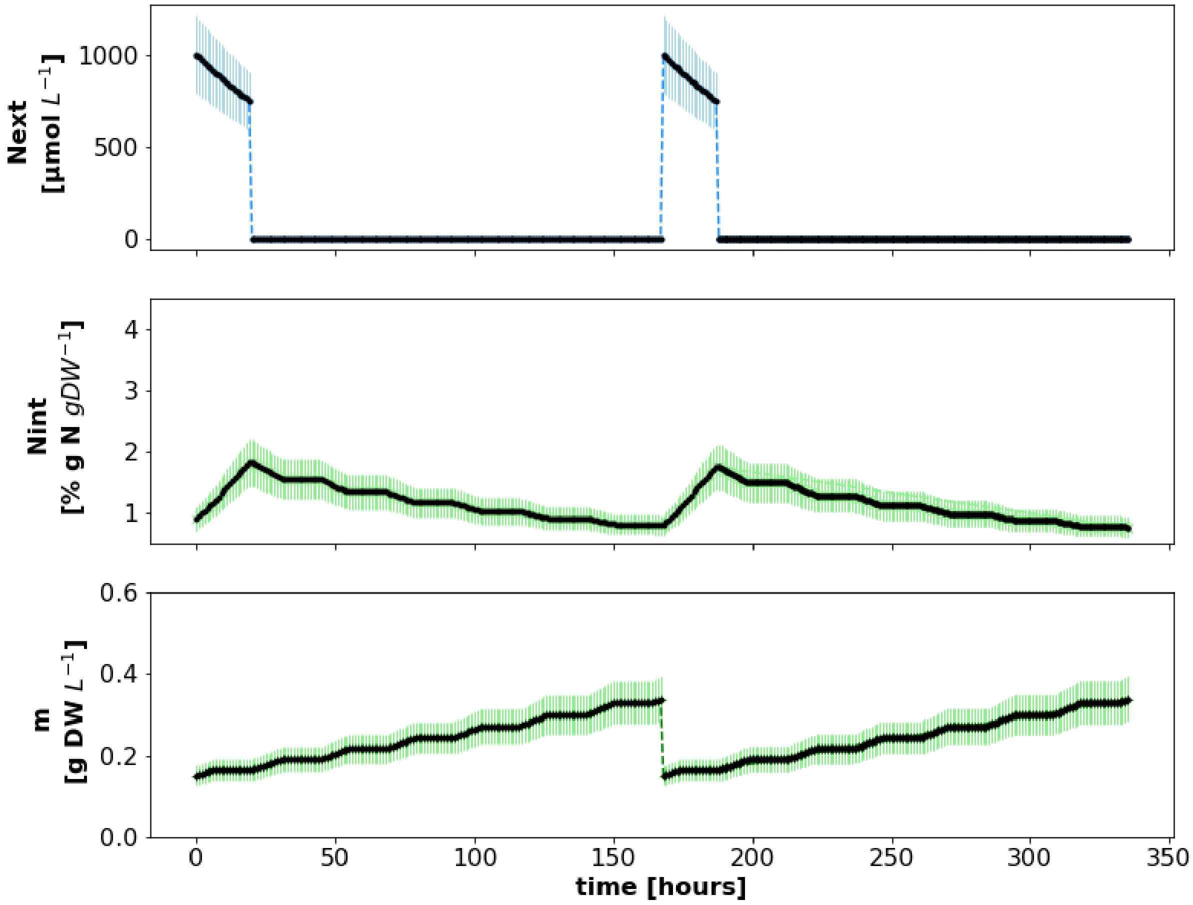


**Figure S14**. Model timewise simulation of Ulva sp. fertilization and cultivation in a controlled photobioreactor. Initial conditions:  Three variables are followed: $N_{ext}$ (µM N), $N_{int}$ (% g N $g^{-1}$ DW), and m (g DW $L^{-1}$). Initial conditions: 0.15 g D.W. per $L^{-1}$ (5 g F.W. per 5 liter reactor), 0.9 % g N $g^{-1}$ D.W. Simulated fertilization included soaking the biomass in 1000 µM N for 20 hours, followed by a week's cultivation under starvation conditions.
